# Chromosome-scale Genome Assemblies of Two Allopolyploid *Cuscuta* Species Uncover Genomic Signatures of Parasitic Lifestyle and Polyploid Evolution

**DOI:** 10.1101/2025.08.18.670985

**Authors:** Tenta Segawa, Shima Yoshizumi, Hiromi Toyonaga, Akira Shiraishi, Kyoko Sato, Takahiro Yamabe, Motoshige Takagi, Masaki Takagawa, Ryusuke Yokoyama, Takehiko Itoh, Eiichiro Ono

**Author notes:** **Corresponding author:** T. Segawa, Research Institute, Suntory Global Innovation Center Ltd., 8-1-1 Seikadai, Seika, Soraku, Kyoto 619-0284, Japan, E. Ono, Research Institute, Suntory Global Innovation Center Ltd., 8-1-1 Seikadai, Seika, Soraku, Kyoto 619-0284, Japan.

## Abstract

Dodders (*Cuscuta* spp.) is an obligated parasitic plant, which lost a large part of photosynthetic genes but gained host genes through parasitism-mediated horizontal gene transfer (HGT). Their migratory ecology across the world would contribute to complexity of the speciation via geographic isolation. Here we report the *de novo* genome assemblies of the two phylogenetically distinct dodders; *C. chinensis* (2n=4x=60) and *C. campestris* (2n=4x=60) that are classified into Clade B and Clade H of subgenus *Grammica*, respectively. Relatively low completeness of BUSCO genes ca.87% indicated progressive gene loss after evolution of parasitic lifestyle due to release from functional constraints as photosynthesis and organ development. Comparative genomics uncovered that both species are the genome size is completely different regardless of the same cytotypes and allopolyploid through the independent ancient hybridization between different parents. Various genomic rearrangements including 1) gene gain and loss events, 2) homoeologous recombination between two subgenomes and 3) the lineage-specific proliferation of transposable elements likely contribute to their genomic diversity and sexual isolation of the two lineages partly sharing their habitats. Our findings not only provide genomic basis to survey parental species for allopolyploidization but also help to understand unique speciation of parasitic dodders through these chromosomal natures.

## Introduction

In plants, polyploidization is one of the major evolutionary drives that generates genetic diversity for morphological and physiological innovation, promotes sexual isolation and environmental adaptation leading to speciation (Leitch and Leitch 2008). Among the different forms of polyploidy, allopolyploidy—arising from the hybridization of genomes from distinct species—is known to involve complex and dynamic changes, including genetic and structural interactions between subgenomes, rewiring of gene expression, chromosomal rearrangements, and reprogramming of epigenetic regulation (Deb et al. 2023, Edger et al. 2017, Hu et al. 2024, Edger et al. 2025). Studies of such allopolyploids are highly relevant to both crop breeding and understanding the mechanisms of plant diversification. However, many questions remain regarding how genome stabilization and functional asymmetry between subgenomes contribute to long-term evolutionary trajectories following polyploidization.

The genus *Cuscuta* (Convolvulaceae in Solanales), parasitic dodder, comprises approximately 200 species of holoparasitic plants and is considered an important model system in plant evolutionary biology due to its extreme morphological reduction and unusual patterns of molecular evolution (Yuncker 1932, Stefanović et al. 2007, Garcia et al. 2014, Costea et al. 2015). Members of this genus lack both roots and leaves, and many species have completely lost the ability of photosynthesis. *Cuscuta* is an obligate parasite that physically connects to the vascular tissues of host plants via haustoria, extracting water, nutrients, and organic compounds including nucleic acids and proteins (Thoday 1911, Kim et al. 2014, Hartenstein et al. 2023). The evolution of such a parasitic lifestyle is accompanied by significant morphological and physiological degeneration and suggests the existence of unique molecular evolutionary trajectories that diverge from those of autotrophic relatives.

Genome assemblies for *Cuscuta* species such as *C. australis* and *C. campestris* have been constructed using next-generation sequencing (NGS) technologies (Vogel et al. 2018, Sun et al. 2018), but they remain at scaffold-level resolution. The previous genomics revealed the unique molecular evolutionary features of the *Cuscuta* lineage as follow; widespread loss of genes involved in photosynthesis, organ development (particularly of leaves and roots), hormonal signaling, and environmental stress responses. Conversely, several studies have demonstrated that *Cuscuta* species acquired genes derived from host plants via horizontal gene transfer (HGT) (Zhang et al. 2014, Yang et al. 2019, Lin et al. 2022). In addition, plastid genome structures vary considerably depending on the degree of photosynthetic capacity, with some species exhibiting extensive gene loss and structural rearrangements (Pan et al. 2023, Park et al. 2019). These loss-and-gain processes highlight the dynamic nature of parasitic plant genomes and provide key insights into their distinct evolutionary trajectories.

Taxonomic classifications have been proposed for the diverse species within the genus *Cuscuta*, largely based on their characters. García et al. (2014) utilized *rbcL* and nuclear large-subunit ribosomal DNA (nrLSU) sequences for phylogenetic analyses and then defined 19 clades (A–S), which they grouped into four subgenera. The subgenus *Grammica* includes clades A through N and contains the largest diversity. *Pachystigma* corresponds to clade P, *Cuscuta* to clades Q and R, and *Monogynella* to clade S, respectively. Among *Grammica*, the section *Cleistogrammica* (clade B) includes *C. campestris*, which has been suspected to be an allopolyploid due to the discrepancies between plastid- and nuclear-derived phylogenies (McNeal et al. 2007, García et al. 2014). However, these analyses failed to elucidate the subgenomic structure potentially resulting from allopolyploidy due to limited sequences in the selected loci.

The genus *Cuscuta* is also known for its remarkable cytogenetic diversity. The ancestral number of chromosomes is estimated to be 2n = 30, which is shared with many *Cuscuta* spp. as well as with related genera such as *Ipomoea* (Ibiapino et al. 2022, Wu et al. 2024). Notably, the number of chromosomes vary widely among species, ranging from 2n = 8 to 2n = 150. This outstanding chromosomal variation suggests progressive polyploidization and genome rearrangements (Pazy and Plitmann 1995). In terms of genome size, *Cuscuta* species also show an enormous variation, with estimates ranging from 270 Mb to over 34 Gb more than a 128-fold difference (McNeal et al. 2007, Ibiapino et al. 2022). Neumann et al. (2021) reported that the genome enlargement is largely attributable to the massive accumulation of transposable elements using short reads only. Due to the unavailability of chromosome-level genome of dodders, chromosomal structural changes via recombination and rearrangement are totally unknown during speciation of dodders.

In this study, we report high-quality genomes of two allopolyploid dodders: *C. campestris*, belonging to subgenus *Grammica*, section *Cleistogrammica* (clade B), and *C. chinensis*, assigned to subgenus Grammica, section *Grammica* (clade H). Both *C. campestris* and *C. chinensis* exhibit similar morphological appearances and partly parasitic preference in terms of their host range, thereby making it difficult to distinguish them based on the morphology in the field. To elucidate the evolutionary nature of allopolyploidy in dodders, we constructed chromosome-scale genome assemblies using PacBio long-read and High-throughput chromosome conformation capture (Hi-C) (Lieberman et al. 2009) sequencing technologies. We further proceeded a series of comparative genomic approaches—including self-alignment analysis, synonymous substitution rate (*d*_S_) estimation, transposable <u>e</u>lement (TE) annotation, and synteny analysis—to explore the evolutionary trajectory of subgenomes, the potential occurrence of recombination between <u>h</u>omoeologous chromosomes (HR), and genome size dynamics. This study would not only help to understand how allopolyploidization shaped the complex genomic architecture in diverged parasitic dodders, but also provide a crucial opportunity to further explore parental and common ancestral species of the allopolyploid dodders prior to hybridization.

## Results

### Karyotyping of C. campestris and C. chinensis

To construct *de novo* reference genomes, we collected individuals of *C. campestris* and *C. chinensis* from two coastal regions in Japan (**Figure 1ab**). Both species were parasitizing *Vitex rotundifolia*, a Lamiaceae species native to the coast. Previous cytogenetic studies reported variation in number of chromosomes for these species, typically ranging from 2n = 56 to 60 (Vogel et al. 2018). Cytological observations showed that both species have the same cytotype of 2n = 60 (**Figure 1cd, Supplementary Fig. S1**). Notably, the chromosome aspect of *C. chinensis* was bigger than that of *C. campestris*. Given that the ancestral basic number of chromosomes in *Cuscuta* are estimated to be 2n = 30 (Ibiapino et al. 2022), these individuals of the different clades are likely tetraploid. We validated the plant material; an individual of each species used for *de novo* genome analysis by genotyping chloroplast genome sequences. We confirmed that both individuals have identical sequences to previously published chloroplast marker sequences of both species (**Supplementary Table S1, S2 and S3**).

**Figure 1.**
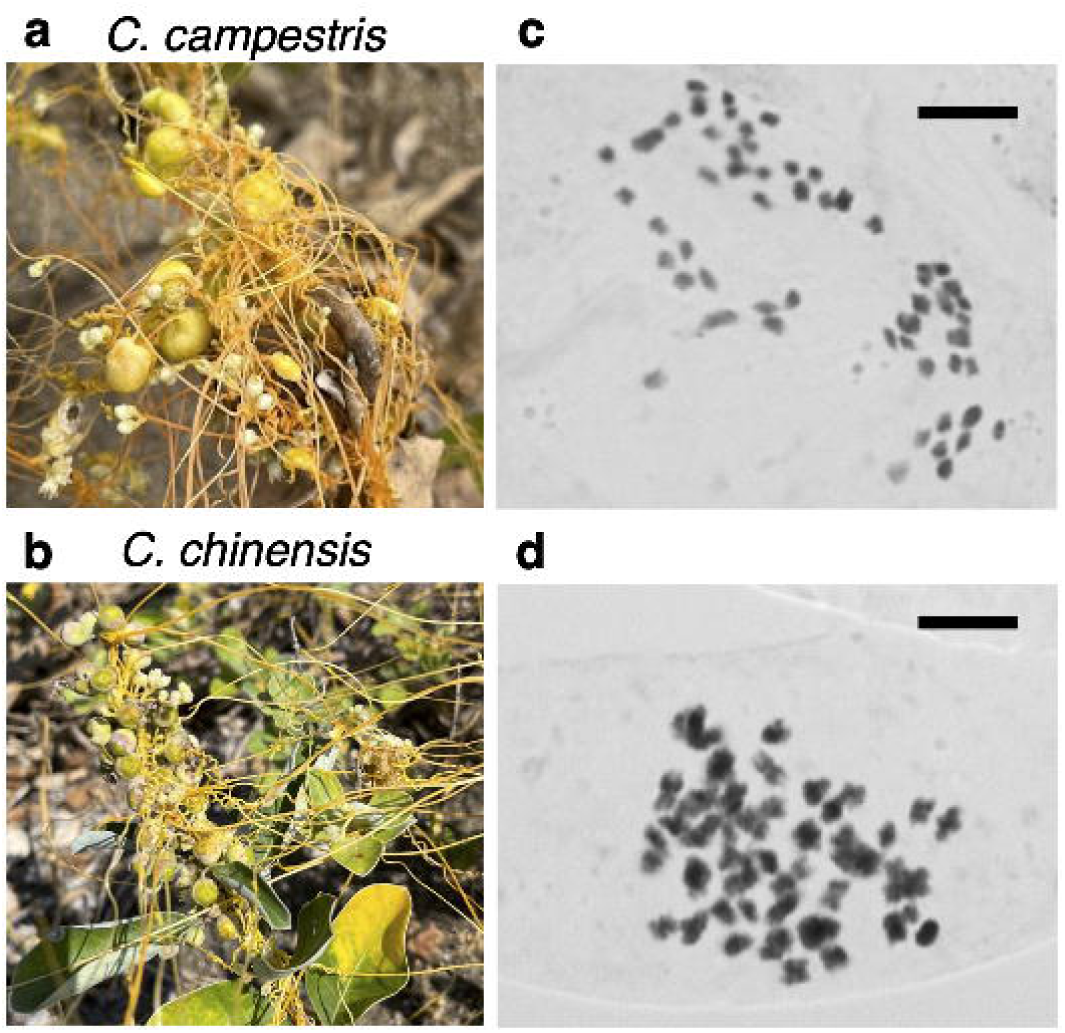
Morphology and chromosomes of C. campestris and C. chinensis. (**a**) and (**b**) show *C. campestris* with insect goals and *C. chinensis*, respectively, parasitizing *Vitex rotundifolia* (Lamiales) in their natural coastal habitats. (**c**) and (**d**) are chromosome photomicrographs of *C. campestris* and *C. chinensis*, respectively. Scale bars represent 5 μm.

### Genome assembly of C. campestris

Using HiFi reads, we constructed a high-quality genome assembly of *C. campestris* with a total length of 514.9 Mb (**Figure 2a, Supplementary Table S4**). This size, estimated to be 493.6 Mb from our k-mer analysis and 611.7 Mb from our flow cytometry analysis, slightly exceeds the previous k-mer-based estimate of 478.7 Mb reported by Vogel et al. (2018), indicating that the assembly is nearly complete (**Supplementary Fig. S2**). The 37 longest contigs accounted for 99.3% of the total genome length, and gene synteny analysis with the phylogenetically related *Ipomoea nil* genome (Hoshino et al. 2016) revealed that each *I. nil* chromosome was covered twice for that of *C. campestris* (**Supplementary Fig. S3**). This suggests that *C. campestris* is an allotetraploid with two distinct subgenome sets. The chromosome numbers were assigned based on homology to that of *I. nil*, and the two subgenome subsets were designated as BI (derived from clade B) and BII (details of this classification are described below). We designated genomic contigs as follows: BI02 corresponds to a single contig homologous to *I. nil* chromosome 2, while some chromosomes such as BI01_1 and BI01_2 are split into two contigs of that to *I. nil* chromosome 1.

**Figure 2.**
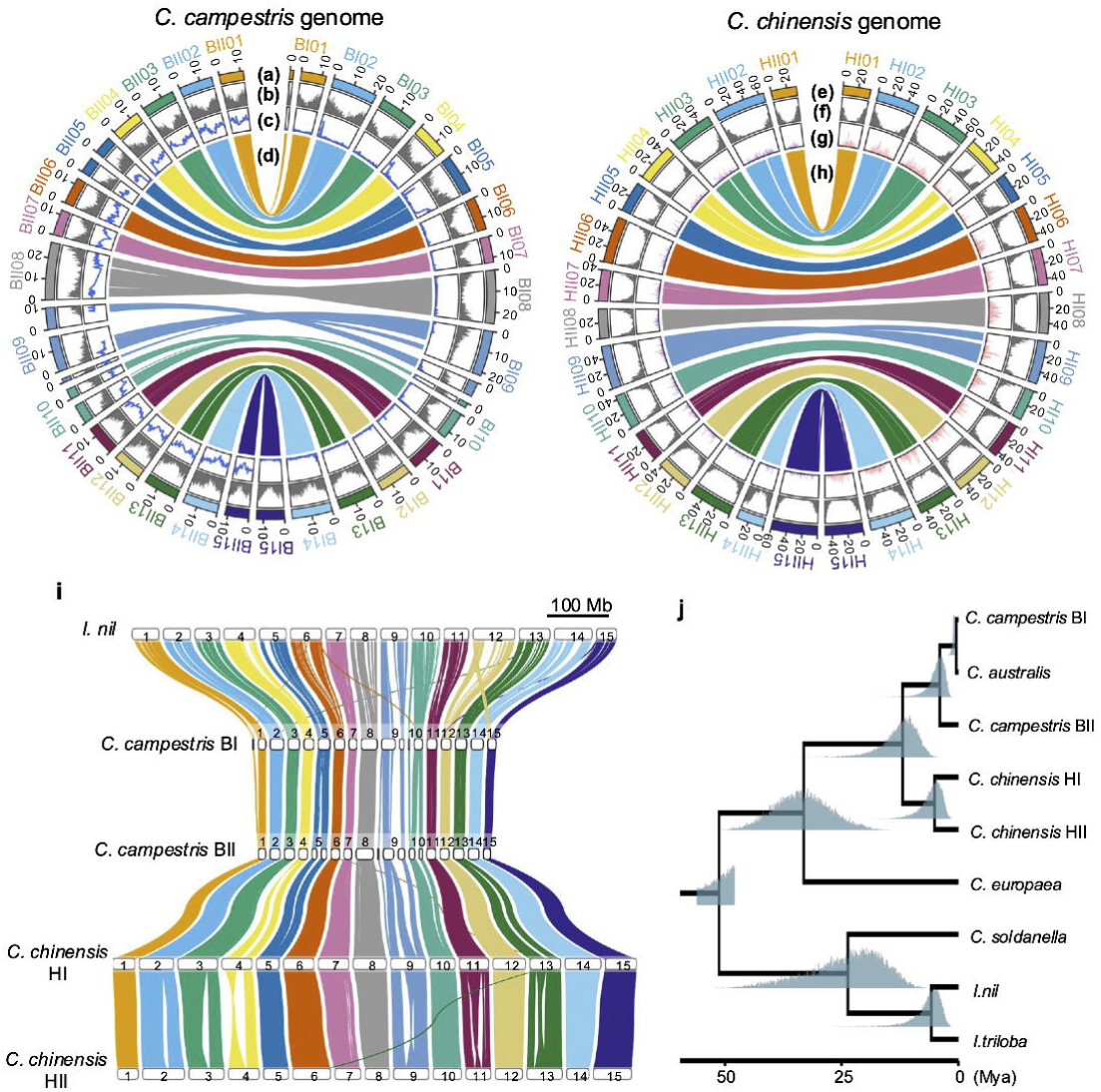
Chromosome-scale allopolyploid genome assemblies of C. campestris and C. chinensis. (**a–d**) Circos plots for *C. campestris*. (**a**) Contig lengths (Mb); (**b**) gene density; (**c**) *d*_S_ between *C. campestris* and *C. australis*; (**d**) syntenic relationships between the two subgenomes. (**e–h**) Circos plots for *C. chinensis*. (**e**) Chromosomes and contig lengths (Mb); (**f**) gene density; (**g**) frequencies of TE families enriched in either the HI or HII subgenomes; (**h**) syntenic relationships between the two subgenomes. (**i**) Comparative genome structure of *I. nil*, *C. campestris*, and *C. chinensis*, showing syntenic regions across corresponding chromosome sets. (**j**) Divergence time estimation within Convolvulaceae. Values at internal nodes indicate the 95% highest posterior density intervals. The blue shaded regions indicate the posterior distributions of node ages inferred from a Markov chain of 20,000,000 iterations. The first 10% of the samples were discarded as burn-in, resulting in 18,000,000 effective samples used for estimation.

From the genome assembly, a total of 37,546 protein-coding genes were predicted, and BUSCO (Benchmarking Universal Single-Copy Orthologs) (Manni et al. 2021) analysis detected 86.7% of complete genes, of which 83.1% were present in more than one copy, suggesting the allopolyploid structure of the genome. Large-scale gene loss across the genome are characteristic features of *Cuscuta*, presumably due to release from functional constraint in parasitic lifestyle. The relative low BUSCO score was comparable to previous reports for *C. campestris* and *C. australis,* 82.1% and 83.7% of BUSCO completeness, respectively (Vogel et al. 2018, Sun et al. 2018). Gene density analysis uncovered reduced gene frequency regions near chromosomal centers, suggesting the presence of centromeric regions (**Fig. 2b**). A centromeric region was identified on each of 30 chromosomes, consistent with the observed haploid number of chromosomes (n = 30). Moreover, telomeric repeats were detected at both ends of 24 chromosomes, indicating that most of the assembly was telomere-to-telomere (T2T) (**Supplementary Table S5**). Collectively, we constructed a high-quality genome assembly covering mono-centromeric chromosomes of *C. campestris* habitant to the Japanese coast.

Previous studies suggested that at least one ancestral genome of *C. campestris* was derived from *C. australis* or its close relative (Costea et al. 2015), but genome-wide certification remained elusive. To elucidate the evolutionary trajectory of subgenomes, we calculated the synonymous substitution rate (*d*_S_) between each *C. campestris* gene and its best homolog in *C. australis*, and compared *d*_S_ distributions across chromosomes. All the chromosomes were clearly classified into two groups: one with a peak at *d*_S_ ≈ 0.00 and the other at *d*_S_ ≈ 0.05 (**Figure 2c**). This bimodal distribution supports the notion that *C. campestris* is an allopolyploid and that one of its subgenomes is closely related to *C. australis*. Therefore, we designated the subgenome closely related to *C. australis* as BI and the other subgenome as BII, and separately referred to them hereafter.

### Genome assembly of C. chinensis

As the *C. chinensis* genome has not been sequenced yet, we constructed the genomic assembly in a similar way to that of *C. campestris* as described above. Using both HiFi and Hi-C reads, the assembly totaled 1.50 Gb, exceeding the *k*-mer-based estimate of 1.22 Gb and being close to the 1.70 Gb estimated by flow cytometry. (**Figure 2e**, **Supplementary Fig. S2, Supplementary Table S4**). The 30 longest contigs accounted for 96.1% of the total assembly, and synteny analysis confirmed that each *I. nil* chromosome was covered twice (**Supplementary Fig. S4**). These results suggest that *C. chinensis* is also an allotetraploid containing two subgenomes as suggested to *C. campestris*. The chromosomes were aligned on *I. nil* chromosomes and then classified into two subgenomes, HI (clade H-derived) and HII (criteria for this classification described below). 37,087 coding genes were predicted, and BUSCO completeness was 87.7% with 74.8% of genes present in two or more copies, comparable to *C. campestris*. Gene density was lower in central chromosomal regions, consistent with typical centromeric localization (**Figure 2f**). Telomeric repeats were identified at both ends of 22 of the 30 chromosomes, showing the high completeness as the T2T assembly (**Supplementary Table S6**).

There were no genome assemblies of the H clade section of *Grammica* comparable to *C. chinensis* and no information of parental lineages for *C. chinensis.* To overcome the unavailability of phylogenetic information of the H clade, we utilized TE composition in each chromosome as a unique genomic signature based on a notion that two subgenomes should independently experience enlargement of different TEs. To infer two subgenomes separately, we selected 27 TE families that showed reliable frequency differences across homoeologous chromosomes and performed principal component analysis (PCA) based on copy numbers. This approach distinguished the highly homologous contigs and classified into two groups, enabling classification into HI and HII subgenomes without parental genomic information (**Fig. 2gh**, **Supplementary Fig. S5**). These results support the notion that *C. chinensis* is not an autopolyploid, but rather an allopolyploid derived from hybridization between two divergent genomes that independently accumulated different TE families in lineage-specific manner.

Comparative genomic analysis between the four *Cuscuta* subgenomes (BI, BII, HI, and HII) and genome of the relative nonparasitic *I. nil* detected several small-scale translocations and rearrangements (**Figure 2i**). For example, part of *I. nil* chromosome 12 showed synteny with chromosome 15 of *Cuscuta*, suggesting a chromosomal rearrangement. While no large-scale chromosomal rearrangements were detected between *C. campestris* and *C. chinensis*, a large inversion was detected specifically on HI04.

Using single copy orthologous gene sets from the assembled genomes, we reconstructed a phylogenetic tree for the Convolvulaceae family including *Cuscuta* spp. (**Figure 2j**). The genus *Cuscuta* diverged from *Ipomoea* and *Calystegia* ca. 50.9 million years ago (Mya) and subsequently split into subgenera *Cuscuta* (*C. europaea*) and *Grammica* (*C. campestris*, *C. chinensis*) ca. 33.3 Mya. Section *Cleistogrammica* (Clade B) and section *Grammica* (clade H) diverged ca. 11.3 Mya, while the progenitor lineages of *C. campestris* and *C. chinensis* diverged more recently at ca. 3.6 Mya and ca. 4.6 Mya, respectively. Furthermore, the BI subgenome of *C. campestris* and *C. australis* were estimated to have diverged around only ca. 0.3 Mya.

### Genomic rearrangements in the allopolyploid genomes

In allopolyploid genomes, recombination between divergent subgenomes—known as homoeologous recombination (HR)—has been frequently reported. To investigate whether HR occurs in *Cuscuta campestris*, we calculated the *d*_S_ between *C. campestris* and *C. australis* genes along each chromosome and surveyed local variation based on the *d*_S_ values between homoeologous chromosome pairs (**Figure 3a**, **Supplementary Fig. S6**). In chromosome BI04, the region from the left end to 3.0 Mb showed *d*_S_ ≈ 0.05, while the downstream region from 3.0Mb showed *d*_S_ ≈ 0.00. In contrast, the corresponding homoeologous chromosome BII04 showed the opposite pattern: *d*_S_ ≈ 0.00 from left end to approximately 3.3 Mb and *d*_S_ ≈ 0.05 downstream from 3.3 Mb. A similar reciprocal shift in *d*_S_ was observed in the regions from the left end to approximately 16.5 Mb and 6.3 Mb of BI05 and BII05_2, respectively. Coverage depth analysis using self-alignment data showed no significant copy number variation in these regions, and allele dosage remained constant at 2 (**Figure 3b, Supplementary Fig. S7**). These results support the occurrence of HR without structural copy number changes. Further analysis identified homoeologous gene pairs— Ccam_Kyo_506–507 on BI04 and Ccam_Kyo_6330–6331 on BII04—where the *d*_S_ shift occurred in opposite directions, indicating a reciprocal pattern of substitution rate change (**Supplementary Fig. S8**). A similar relationship was observed between gene coding regions Ccam_Kyo_28037–13413 on BI05 and Ccam_Kyo_24229–9481 on BII05. These patterns indicate that reciprocal HR events frequently occurred between subgenomes after hybridization between parental dodders (**Figure 3c**).

**Figure 3.**
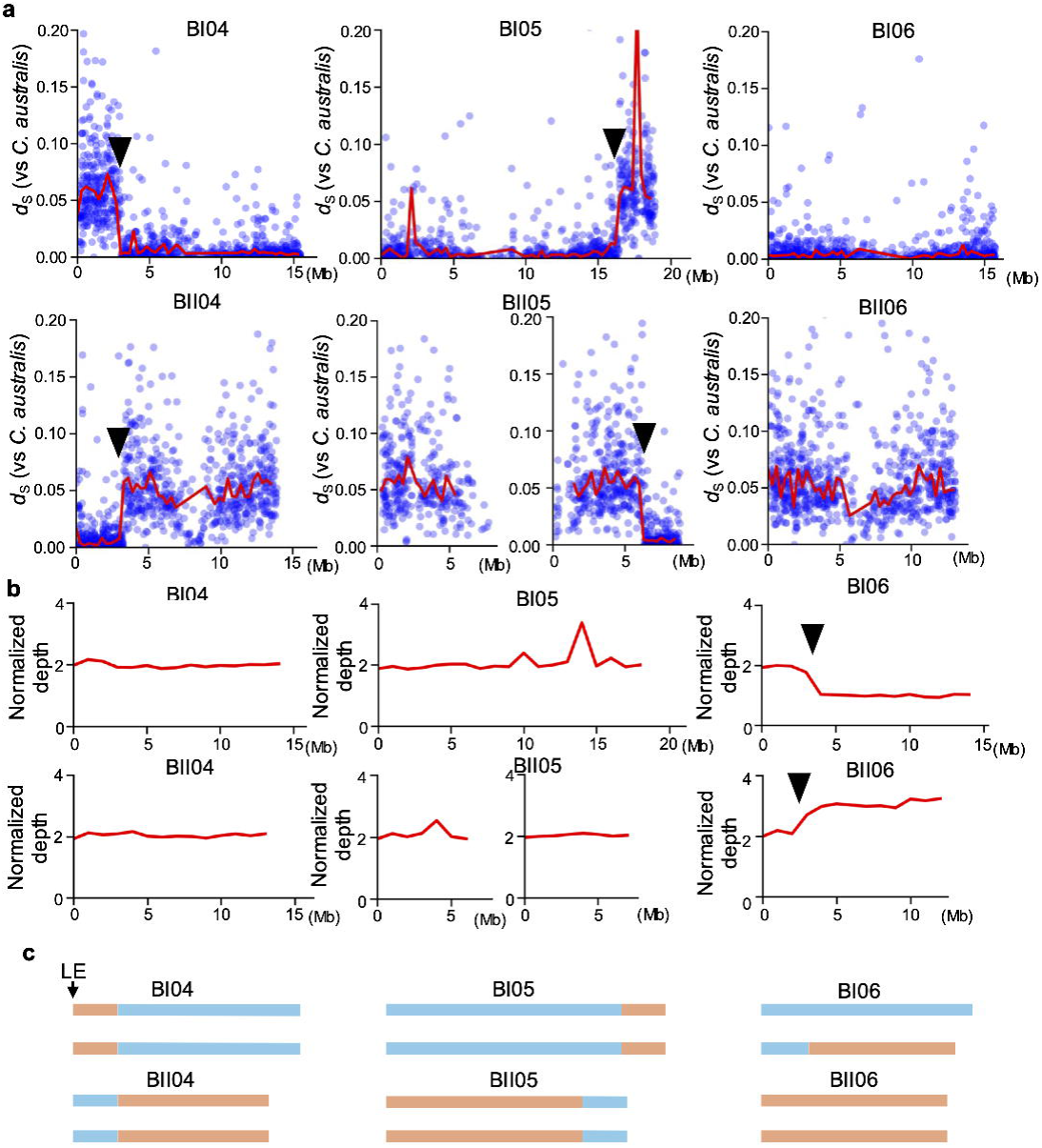
Homoeologous recombination (HR) in C. campestris. Reciprocal HR were detected on BI04-BII04 and BI05-BII05, while non-reciprocal HR was identified on BI06-BII06 in the *C. campestris* genome. (**a**) The blue plots show the *d*_S_ values with *C. australis* genes at each gene position. The red lines indicate the median *d*_S_ value between *C. campestris* and *C. australis* of 300 kb bins at each chromosomal position. Black arrowheads indicate reciprocal HR sites. (**b**) The red line shows the normalized self-aligned depth of 1Mb bins in each chromosome position. Normalized depth is a representation of genome dosage. Black arrowheads indicate non-reciprocal HR sites. (**c**) Predicted chromosome structure of the *C. campestris* genome, inferred from *d*_S_ values and normalized depth. Orange regions represent chromosomes derived from an ancestral species more closely related to *C. australis*, while light blue regions represent chromosomes derived from another ancestral species. LE: Left end.

To further explore the possibility of non-reciprocal HR in *C. campestris*, we compared coverage depth between BI06 and BII06 (**Figure 3b, Supplementary Fig. S7**). In BI06, the region from the left end to approximately 3.8 Mb had normalized depth to be 2, which dropped to 1 beyond this point. Conversely, in BII06, normalized depth was 2 from left end to approximately 3.2 Mb but thereafter increased up to 3 (**Supplementary Fig. S9**). This pattern suggests that one allele in BI06 was retained while the other was replaced by a segment from BII06, indicative of a non-reciprocal HR. The resulting allele frequency is locally biased to BI06:BII06 = 1:3 at this region (**Figure 3c**).

In *C. chinensis*, for unavailability of genomic information of the ancestral plants, we inferred non-reciprocal HR by calculating *d*_S_ distributions and chromosome structure instead of the direct comparison of *d*_S_ values with reference genome of the ancestral species. Coverage depth comparisons between HI and HII chromosomes showed no significant allelic bias, suggesting that both subgenomes are stably conserved in equal ratio (**Supplementary Fig. S10**). However, *d*_S_ survey across HII chromosomes showed some restricted regions, particularly at chromosome ends, with *d*_S_ ≈ 0.00, contrasting with *d*_S_ ≈ 0.05 in most other regions (**Supplementary Fig. S11**). Further detailed analysis indicated that these low-*d*_S_ regions—HII07 (around 39.0 Mb), HII08 (around 38.3 Mb), HII10 (around 0.2 Mb), and HII13 (around 0.4 Mb)—are likely the result of non-reciprocal HR. These segmental non-reciprocal HR (snrHR) were observed as a 4:0 coverage ratio in which one ancestral haplotype is completely replaced by another. These results support the notion that these subgenomes in allopolyploids are separately conserved but occasionally experienced reciprocal, non-reciprocal HR and snrHR. These chromosomal rearrangements would contribute to increasing genetic diversity.

### Evaluation of transposable elements in two dodder genomes

The subgenomes of *C. chinensis* (HI and HII) are approximately 2.6–3 times larger than those of *C. campestris* (BI and BII) despite the same cytotype with that of *C. campestris* **(Figure 1cd)**. To clarify the molecular insights of the genomic enlargement, we surveyed TEs across the entire genomes using RepeatMasker. As a result, the total length of repetitive elements in *C. chinensis* was approximately 3.9 times greater than that in *C. campestris*, showing a clear positive correlation between TE accumulation and genomic enlargement (**Figure 4a**). In contrast, the size of non-repetitive regions was comparable between the two species, with each subgenome comprising approximately 91.8–97.0 Mb. Collectively, these results show that the difference in genome size of the two dodders is primarily attributable to the enlargement of repetitive elements, in which, LTR retrotransposons were the predominant family.

**Figure 4.**
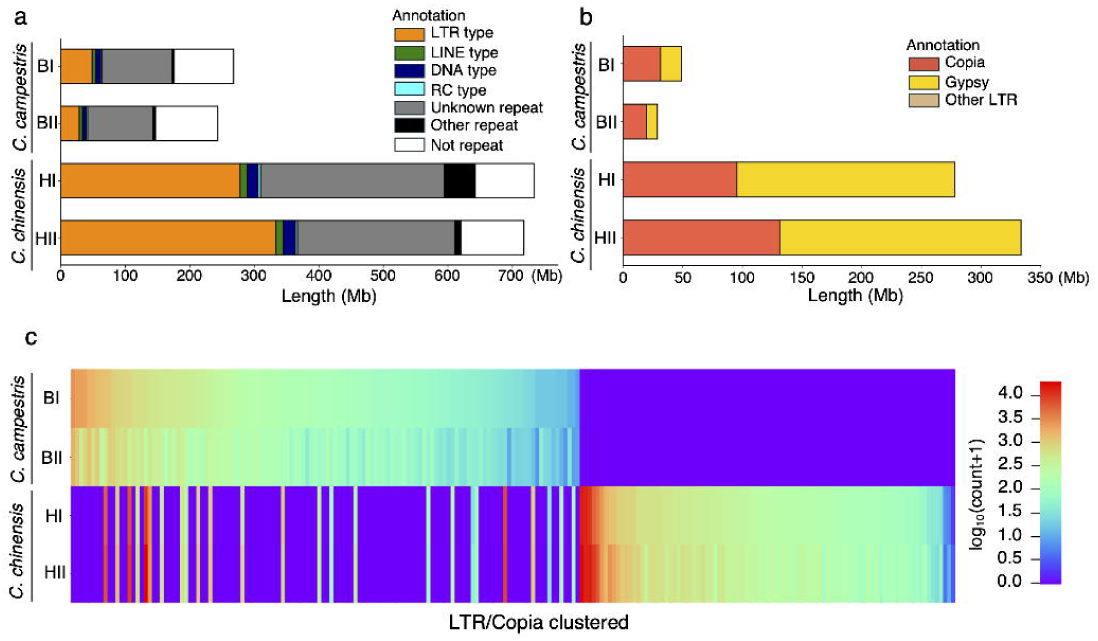
Composition and comparison of TE. (**a**) and (**b**) show stacked bar plots representing the total length of repeat elements in each subgenome of *C*. *campestris* and *C*. *chinensis*. (**a**) Repeat elements annotated by RepeatMasker. Unannotated regions are shown in white. (**b**) Detailed annotation of LTR-type TEs only. (**c**) Heatmap showing the count of LTR/Copia elements clustered based on ≥80% sequence identity.

Among various repetitive elements, Copia- and Gypsy-type LTR elements appeared to be the major TE classes, in *C. chinensis* genome exhibiting a higher proportion of Gypsy elements than *C. campestris* (**Figure 4b**). While Copia-type was abundant in both species, different Copia clusters were observed in each lineage (**Figure 4c**), suggesting that lineage-specific changes of Copia-type TE occurred after the divergence of the two sections. We further assessed temporal change of TE using the Kimura substitution level of each LTR family and found that both species similarly exhibited a peak at a Kimura substitution level around 5, despite the different composition of the TEs. Notably, in *C. chinensis*, Gypsy and Copia elements seem to be differentially amplificated (**Supplementary Fig. S12**). Furthermore, few TEs clearly inherited from a common ancestor were found in the present genomes of the two species **(Figure 4c)**, indicating that their present genome size had been mainly influenced by recent transposon activity rather than ancient genome size. Taken together, these results indicate that 1) lineage-specific LTR proliferation and reduction occurred independently after the divergence of section *Cleistogrammica* and section *Grammica*, 2) the recent TE activity would have contributed to the present genome size of *C. chinensis* and *C. campestris*.

### Origin of C. campestris

Based on the phylogenetic classification proposed by Costea et al. (2015), section *Cleistogrammica* can be further subdivided into three lineages, which we refer to here as BI, BII, and BIII for clarity. BI includes *C. australis* and *C. obtusiflora*; BII includes *C. glabrior* and *C. runyonii*; and BIII includes *C. pentagona* and *C. harperi*. To investigate the maternal lineage of *C. campestris*, we extracted the chloroplast genome from the raw assembled *C. campestris* contigs and performed BLAST searches using *rbcL* and *trnL-F* sequences as previous phylogenetic analyses (Garcia et al. 2014, Costea et al. 2015) (**Supplementary Table S1 and S2**). Both sequences matched previously published *C. campestris* sequences with 100% identity, and among the three clades, they showed the highest similarity to sequences from the BII clade. This result is consistent with the previous phylogenetic studies and suggests that a BII-clade species served as the maternal progenitor of *C. campestris*.

Costea et al. (2015) also proposed that the paternal lineage of *C. campestris* may be derived from a hybridization event involving BI and BIII clade species. To test this hypothesis, we compared RNA-seq data of *C. pentagona* (BIII clade) based on *d*_S_ values between *C. pentagona* and each subgenome (BI and BII) of *C. campestris* (**Supplementary Fig. S14**). If *C. pentagona* were involved in the ancestry of *C. campestris*, we would expect to observe regions in the BI subgenome where *d*_S_ values relative to *C. pentagona* are lower than those relative to *C. australis*. However, no such regions were observed when reciprocal HR regions were properly considered. These results suggest that the BI subgenome of *C. campestris* is basically derived from the BI clade including *C. australis* and its close relatives. In our investigation, the BII clade likely contributed to the maternal genome based on genomic similarity as previously predicted (Costea et al 2014). This result was further confirmed by the chloroplast genome sequence. For this reason, we concluded that the BI clade contributed to the paternal genome. No evidence of genomic contribution from the BIII clade was found in the birth of *C. campestris*, however we did not rule out the possibility of a different type of allopolyploids in *C. campestris* which are widespread and geographically isolated in the world.

### Gene loss and gain in dodder genomes

To seek gene loss and gain in the two dodders, we compared ortholog groups (OG) identified by OrthoFinder among the newly assembled genomes of *C. campestris* and *C. chinensis*, along with those of *C. europaea* (GCA_945859875.1), *Ipomoea nil* (Hoshino et al. 2016, Asagao_1.2 genome; http://viewer.shigen.info/asagao/index.php), *I. triloba* (GCF_003576645.1), and *Solanum lycopersicum* (GCF_000188115.5) in Solanales. We identified 2,067 OGs that were commonly present in the non-*Cuscuta* species (*Ipomoea* and *Solanum*) but specifically absent across all *Cuscuta* genomes. On the other hand, we found 315 OGs that were specifically shared among all the *Cuscuta* species but not found in the non-parasitic relative species.

Gene Ontology (GO) analysis showed reduced-67 GOs and enriched-226 GOs in dodder genomes, respectively (**Supplementary Table S7 and S8**). We found a reduction for biological processes related to photosynthesis (e.g., photosystem I/II assembly, response to high light intensity), carbohydrate metabolism (e.g., starch and fructose catabolism), and oxidative stress responses in dodders (e.g., glutathione peroxidase activity and detoxification pathways) (**Supplementary Fig. S15**, **Supplementary Table. S9**). Genes involved in plant-type cell wall organization and hormone signaling—such as those responsive to abscisic acid and salicylic acid—were also highlighted as the reduced genes. These results are consistent with the reduction of self-sustaining functions in *Cuscuta* as previously suggested (Sun et al., 2018; Vogel et al., 2018), reflecting its obligate parasitic lifestyle. Moreover, GOs associated with ceramide catabolism, lignan biosynthesis and nitrate transport were also reduced in dodder genomes, indicating that numerous metabolic alterations occurred after divergence from autotropic relatives.

In contrast, gene groups were enriched in functions related to interactions with other organisms, including terms such as “modulation of process of another organism” and “toxin activity” (**Supplementary Fig. S16**, **Supplementary Table. S10**). These gene gains also included proteases (e.g., cysteine- and aspartic-type peptidases), beta-glucosidases, and nutrient reservoir-related proteins, which may play roles in host cell wall degradation, nutrient uptake, and storage. We also found the GOs associated with RNA regulation, phytochrome B and plasmodesmata are enriched. Taken together, the dynamic genetic alteration likely associated with parasitism in *Cuscuta*, namely the loss of genes potentially involved in autotrophic and independent metabolic processes, and the gain of genes potentially involved in host manipulation and resource acquisition.

## Discussion

In this study, we showed that both *C. chinensis* and *C. campestris* are allopolyploid with the same cytotype but quite different genome sizes, which is largely influenced by recent TE activity. Multiple genetic processes—including allopolyploidization, chromosomal rearrangements between subgenomes, gene gain and loss and lineage-specific TE proliferation and reduction—have shaped the complicated speciation in *Cuscuta*. Previous studies implied the entangled evolutionary trajectories of dodders through the allopolyplodization (Garcia et al. 2014, Costea et al. 2015). Furthermore, accelerated mutation rate would further increase complexity of the genomic evolution (Bromham et al. 2013). However, our data suggests that the frequency of chromosomal recombination between the subgenomes is basically rare.

HR is widely reported in allopolyploid species such as *Brassica napus* (Higgins et al. 2021) and hexaploid bread wheat (Zhang et al. 2020). Allopolyplodization is thought to provide opportunities for HR, thereby increasing chromosomal complexity and chance of adaptation. Genome-wide scanning of *d*_S_ values and allele dosage indicated multiple reciprocal and non-reciprocal HR events across several chromosomes between subgenomes. Important to note, non-reciprocal HR between the BI06 and BII06 chromosomes was maintained heterozygously, suggesting a selection for this region of both subgenomes, although distribution of the HRs should be further surveyed throughout populations. Our findings support the notion that HR contributes to increasing genomic diversity (Edger et al. 2025), thereby occasionally acquiring adaptive traits.

In *C. campestris*, subgenome analysis revealed that the BI subgenome likely originated from a lineage closely related to *C. australis*. Although the genome of the true progenitor corresponding to the BII subgenome is currently unavailable, chloroplast genome analysis suggested the maternal genome is derived from clade BII species such as *C. glabrior* or *C. runyonii* based on the highest similarity of the sequences. These findings herein are basically consistent with the intraclade classification of section *Cleistogrammica* proposed by Costea et al. (2015) and provide strong genomic evidence that *C. campestris* is an allotetraploid derived from two separate ancestral species. In contrast, no support was found for a contribution from clade BIII (*C. pentagona*), which was previously hypothesized as a potential progenitor. This suggests the possibility of parallel allopolyploidizations in geographically isolated populations of this species.

*C. chinensis* and *C. campestris* are two phylogenetically distinct allopolyploid dodders with the same cytotype; 2n=60 chromosome numbers. It should be noted that there are outstanding variations in the basic chromosome numbers of the two allopolyploid dodders, e.g., 2n=16, 28, 15II, 56, and 60 were reported for *C. chinensis* (Asano 1960, Sampathkumar 1979, Aryavand 1987, Měsĺček and Soják 1995, Vasudevan 1975), while n=14, 28, 2n=28, 56, c. 56, and 60 for *C. campestris* (Finn and Safijovska 1933, Finn 1937, Fogelberg 1938, Ward 1984, Aryavand 1987, García and Castroviejo 2002, Vogel et al. 2018, García et al. 2019, Taşar et al. 2022, Ibiapino et al. 2022). Thus, it is doubtful that multiple species are included into the same species name and miscalculation of chromosome numbers. Thus, further multiple sampling of these dodders from geographically diverse populations followed by cytogenetic and genomic analysis is needed to clarify the controversial cytogenetic and phylogenetic issues.

We found that genome size of the HI and HII subgenomes of *C. chinensis* (1.50 Gbp) are approximately 2.6–3 times larger than those of *C. campestris* (514.9 Mbp) despite the same cytotypes, and this size difference was shown—at the whole-genome level for the first time—to be primarily attributed by lineage-specific accumulation and reduction of TEs. Such amplification of LTR elements is consistent with previous report by Sun et al. (2018), who suggested that elevated genome size in *Cuscuta* species is largely attributed by the accumulation of transposable elements. Moreover, such differential trajectories of TE after speciation may have contributed to section-specific diversification. Important to note, the TE configuration is a unique genomic signature maintained in each of the subgenomes after hybridization, by which we could split the allopolyploid genomes into two subgenomes without parental references. This approach would be widely applicable to other orphan allopolyploids like *C. chinensis* unless chromosomal recombination is highly active.

*Cuscuta* species have undergone substantial gene loss, particularly in pathways related to photosynthesis, carbohydrate metabolism, oxidative stress response, and hormone signaling. Low completeness of BUSCO genes ca. 87% in the genomes of two dodders would support the idea that many genes are lost from the genome of the most recent common ancestral parasitic dodder due to their unnecessity for parasitic lifestyle. A large part of gene losses would be tightly associated with their extreme morphological simplification and obligate dependence on host plants, consistent with previous genome analyses of *Cuscuta* (Sun et al. 2018, Vogel et al. 2018).

In contrast, we found gene families uniquely present in parasitic dodders but absent in non-parasitic relatives, many of which are likely associated with host interacting functions. Of note, the enriched GOs in this analysis are expected to include both genes of endogenous gene amplification and exogenous horizontal gene transfer (HGT) derived from host plants (Yang et al., 2019), the former of which likely contribute to parasitism evolution but latter of which likely contribute to reinforcement of parasitism since the ability of parasitism is prerequisite for HGT. It is remarkable that GOs associated with cell wall modulation are obviously enriched in dodders (**Supplementary Table S10**). Considering that some of the enzymes in the GOs are secreted by haustoria, the enriched genes for cell wall remodeling might be directly associated with penetration process by degrading host tissues and facilitating nutrient uptake via the haustorium. Furthermore, the enriched GOs associated with phytochrome B (PHYB) and plasmodesmata might accord with a recent work on PHYB2 involved in light-mediated host recognition and parasitism upon the haustorium formation in the early stage of parasitism (Yokoyama et al. 2025) and previous works on interspecific plasmodesmata involved in cytoplasmic translocation of various RNA molecules between host and parasitic plants (Fischer et al. 2021, Li et al. 2022).

A schematic summary of the genomic evolution of *C. campestris* and *C. chinensis* is shown in **Figure 5**. Our data provides a framework for understanding how allopolyploids have undergone genomic specialization. Importantly, the pattern of gene loss and gain would be partly comparable to that of other parasitic plants, such as *Striga* and *Orobanche*. These species (Lamiales) are phylogenetically distant to dodders (Solanales) but similarly reduced photosynthetic gene repertoires and increased gene sets related to host-parasite interface formation and nutrient acquisition (Yang et al. 2019). The parallel evolution of such genomic traits among phylogenetically unrelated parasitic plants suggests common selective pressures and adaptive constraint for their unique parasitic lifestyle. Notably, the common genetic changes observed in different parasitic plants would be based on the precedent parasitic ability, however the origin and evolution of parasitic ability remains unknown, basically due to the technical difficulty of genetic approaches on most obligate parasites. Temporal divergence analysis of the parasite genomes in comparison with non-parasitic relatives might provide a clue to assess early genetic changes for parasitic ability.

**Figure 5.**
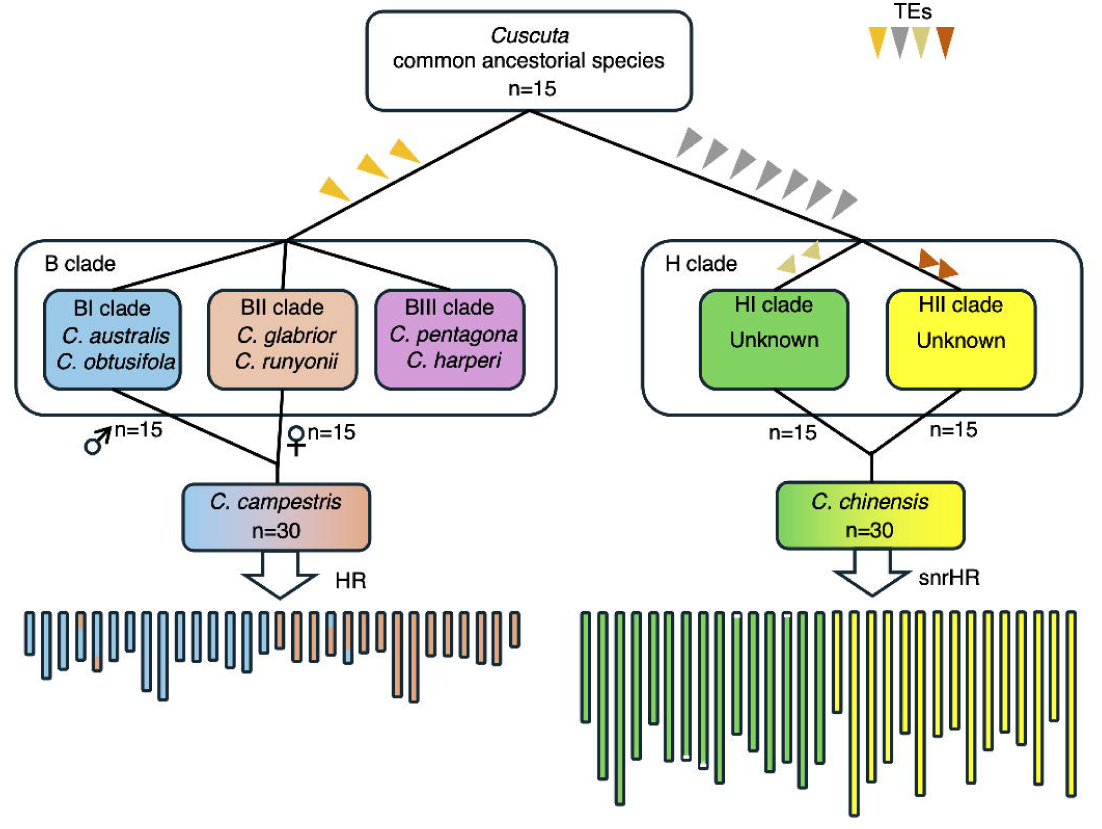
Summary of genomic evolutionary trajectories in C. campestris and C. chinensis. Schematic representation of genome evolution events revealed by genome assemblies of *C. campestris* and *C. chinensis*. The upper panel shows a phylogenetic tree depicting divergence from the common ancestor of *Cuscuta* and *Ipomoea*, followed by species diversification and allopolyploidization events. Insertion events of repeat elements are indicated by triangles on the corresponding branches. The lower panel illustrates genome structures of the allopolyploids, including homoeologous recombination between subgenomes. HR: Homoeologous Recombination. snrHR: segmental non-reciprocal Homoeologous Recombination.

## Materials and Methods

### Plant materials and Next-Generation Sequencing

Mature seeds of *C. campestris* were collected at the coast of Sea of Japan in Kumihama-cho, Kyotango, Kyoto Prefecture, Japan. After germination in a greenhouse, seedlings were parasitized onto *Sesamum indicum*, and a single individual was harvested for genome sequencing. Aerial tissues of *C. chinensis* were collected at coast of Pacific Ocean in Minami-cho, Kaifu, Tokushima Prefecture, Japan These plants were propagated by parasitizing *Nicotiana benthamiana* in a greenhouse at Tohoku University. Genomic DNA (gDNA) from a single individual of both species was extracted using the NucleoBond HMW DNA (MACHEREY-NAGEL) following the manufacturer’s protocol. High-quality gDNA was subjected to whole-genome sequencing using the PacBio Revio platform to obtain HiFi reads (**Supplementary Table S11**). For RNA-seq data used in gene prediction, fresh aerial tissues of *C. chinensis* parasitic individuals were collected and total RNA was extracted with RNeasy Plant Mini Kit for RNA Extraction (QIAGEN) or ReliaPrep RNA Tissue Miniprep system (Promega). Strand-specific libraries were constructed after polyA selection using NEB Next Poly(A) mRNA Magnetic Isolation Module and NEB Next Ultra II Directional RNA Library Prep Kit, and 150 bp paired-end (PE) sequencing was performed on the Illumina NovaSeq X platform (**Supplementary Table S12**). The Hi-C library for *C. chinensis* was constructed using the Dovetail Omni-C Kit (Dovetail Genomics, Scotts Valley, CA, USA) according to the manufacturer’s protocol and was subsequently sequenced on a NovaSeq X Plus platform (**Supplementary Table S13**).

### Chromosome observation

Samples used for chromosome observation were derived from seeds of the same wild populations collected at the respective sampling sites. For chromosome counting, shoot apices from germinated seedlings were pretreated in 2.2 mM 8-hydroxyquinoline at room temperature (approximately 25°C) for 1 hour and subsequently kept at 5°C for 15 hours. After pretreatment, root tips were fixed in a 1:3 mixture of glacial acetic acid and absolute ethanol for 1 hour at room temperature, followed by maceration in 1 M hydrochloric acid at room temperature for 4 hours and at 60°C for 10 minutes. The tissues were then rinsed in running water, stained with 1.5% lacto-propionic orcein, squashed, and observed under a light microscope to find cells during metaphase.

### Genome size estimation using flow cytometer and k-mer analysis

For genome size estimation by flow cytometry, nuclei were isolated from *A. thaliana* and Cuscuta using a razor blade to chop tissues in a buffer prepared by 4-fold dilution of the Nuclei Isolation Buffer 4× (CelLytic™ PN Isolation/Extraction Kit, Sigma-Aldrich) containing Hoechst 33342 Staining Dye Solution (Abcam) at a 1:4000 dilution. The chopped samples were incubated in the dark at 4 °C for 10 min for nuclear staining, and filtered through a 35 μm mesh. The prepared samples were analyzed with a BD FACSMelody™ flow cytometer (BD Biosciences) using UV excitation. To estimate genome size, *k*-mer frequency analysis was performed using HiFi reads. First, Jellyfish ver 2.2.10 (Marçais and Kingsford 2011) was used to count the occurrences of 32-mers with the following parameters: -m 32 -s 1G -C. The resulting *k*-mer count database was then converted into histogram format using the jellyfish histo command. The generated *k*-mer histogram was analyzed using GenomeScope2 ver 2.0.1 (Ranallo-Benavidez et al. 2020), with the *k*-mer size set to 32 for genome size estimation.

### Genome assembly

The *C. campestris* genome was assembled using HiFi reads with Hifiasm ver 0.25.0 (Cheng et al. 2021). The resulting primary contigs were aligned, together with the alternative contigs, to the *Ipomoea nil* reference genome using minimap2 ver 2.28 (with “-x asm20” option) (Li 2021) to identify regions that were not covered by either of the two subgenomes. These regions, uncovered in the primary contigs, were then supplemented using the corresponding sequences from the alternative contigs to construct an integrated assembly. To remove organellar sequences, the assembled contigs were aligned using minimap2 ver 2.28 (with “-x asm20” option) against known mitochondrial (BK016277.1) and chloroplast (NC_052920.1) genome sequences. Contigs that matched these organellar genomes were excluded from the final assembly. The *C. chinensis* genome was assembled using HiFi and Omni-C reads with Hifiasm ver 0.25.0 in Hi-C mode. The resulting primary contigs (p_ctgs) were scaffolded with YaHS v1.2 (Zhou et al. 2023) using the Omni-C data and the parameters “-q 10 --no-contig-ec”. The Hi-C contact map was visualized with Juicebox v1.11.0828 (Robinson et al. 2018), and manual inspection was conducted to assess potential mis-assemblies, telomeric repeat positions, and scaffold orientation. Based on this inspection, only one likely missing join between contigs was manually corrected, and no mis-assemblies or incorrect orientations were detected.

### Repeat annotation and gene prediction

The assembled genome sequences were used to construct a custom repeat element database with RepeatModeler ver 2.0.6 (Flynn et al. 2020). Repeat annotation and masking were performed using RepeatMasker ver 4.1.7, generating both hard-masked and soft-masked versions of the genome. These masked sequences were then used as inputs for gene prediction with Ginger ver 1.0.1 (Taniguchi et al. 2023) and BRAKER3 ver 3.0.8 (Gabriel et al. 2024), respectively. For Ginger-based prediction, RNA-seq data listed in **Supplementary Tables S12 and S14** were used, with protein sequences from Ipomoea triloba and *I. nil* specified as reference proteins. The SPALNDB parameter was set to “Angiosp”, and other parameters were kept at default values. For BRAKER3, RNA-seq reads were first merged and aligned to the assembled genome using HISAT2 ver 2.2.1 (Kim et al. 2019). The resulting SAM files were converted to BAM format with SAMtools (Li et al. 2009) and used as input. Protein sequences from the OrthoDB11 Viridiplantae dataset were specified as the reference, and BRAKER3 was executed with the --softmasking option enabled. The outputs from Ginger and BRAKER3 were integrated by the following procedure: Genes not predicted by Ginger or identified as partial/incomplete were replaced with predictions from BRAKER3. Finally, the predicted gene models were annotated by comparing them against the NCBI nr database using DIAMOND blastp ver 2.1.11 (Buchfink et al. 2021). The top hit annotations were retained unless the hit was labeled as “uncharacterized protein”, “unnamed protein”, or “hypothetical protein”, in which case the gene was designated as “uncharacterized protein”.

### Synteny, synonymous substitution distance and coverage depth

Synteny analysis was performed by first conducting pairwise similarity searches of protein sequences using DIAMOND blastp with an e-value cutoff of 1e-10. The resulting alignment files were used as input for MCScanX (Wang et al. 2012) to identify syntenic blocks. For comparisons involving homoeologous genes in allopolyploid genomes, the --max-target-seqs 2 option was used in DIAMOND to allow for detection of duplicated orthologs. In all other comparisons, --max-target-seqs 1 was applied to restrict results to the top hit. To calculate *d*_S_, DIAMOND blastp was performed between target protein sequences with an e-value threshold of 1e-20.

Homologous pairs in which ≥90% of the query sequence was aligned were extracted using a custom script. The identified homologous protein pairs were aligned using MAFFT ver 7.525 (--auto option) (Katoh 2005), and codon alignments were generated with PAL2NAL ver 14.1 (Suyama and Bork 2006). The resulting codon alignments were then used as input for codeml (yn00 method) in the PAML package ver 4.10.7 (Yang and Nielsen 2000, Yang 2007) to estimate *d*_S_ values.

HiFi reads were aligned to the assembled genome using minimap2 with the -ax map-hifi option. The resulting SAM files were converted to BAM format using SAMtools. Coverage depth across defined bin regions was calculated using the SAMtools coverage command. Normalized coverage depth was computed using the following formula: Normalized depth = (coverage depth of each bin) × 2 / (median coverage depth across all bins)

### Divergence time estimation

Divergence time estimation was performed using the assembled subgenomes of *C. campestris* (BI and BII) and *C. chinensis* (HI and HII), along with genome data from *Cuscuta australis* (GCA_945859875.1) (Sun et al. 2018), *C. europaea* (GCA_945859875.1) (Neumann et al., 2023), *Ipomoea nil*, *I. triloba* (GCF_003576645.1) (Wu et al., 2018), *Calystegia soldanella* (GCA_964235135.1), *Capsicum annuum* (Zunla-1 v2.0: http://www.bioinformaticslab.cn/files/genomes/pepper), *Solanum lycopersicum* (GCF_000188115.5), *Nicotiana attenuata* (GCA_001879085.1) and *Arabidopsis thaliana* (GCA_000001735.1) (Lamesch et al. 2012). Orthofinder (Emms and Kelly 2019) analysis was conducted using the 524 single-copy orthologous genes shared among all taxa were identified. These gene sequences were aligned using MAFFT (--maxiterate 1000 --localpair), and codon alignments were generated with PAL2NAL. A total of 6,883 fourfold degenerate third-codon transversion (4DTv) sites were extracted from codon alignments, and divergence time estimation was conducted using BEAST v2.7.7 (Bouckaert et al. 2019). BEAUti was configured with the following parameters: Substitution Rate = “Tree” (estimated); Gamma Category Count = 4; Shape = estimated; Substitution Model = GTR; Clock Model = Relaxed Clock Log Normal; MCMC Chain Length = 20,000,000. The calibration points were set as described below: The divergence between *A. thaliana* (Brassicales) and Solanales: 85.80–128.63 Mya (Morris et al. 2018); The divergence between Solanaceae and Convolvulaceae: 59.1–83.9 Mya (Eserman et al. 2014); The divergence between *Cuscuta* and other Convolvulaceae (*Ipomoea* and *Calystegia*): 47.8–56.0 Mya (Muller 1981).

### GO enrichment analysis

Predicted protein sequences were first searched against the NCBI nr database using DIAMOND blastp (-f 5) with the output format set to XML. In parallel, functional domains were annotated using InterProScan (-f xml --disable-precalc) (Jones et al. 2014) to generate XML-formatted annotation results. The two XML files were then merged and processed in Blast2GO (Conesa et al. 2005) to assign GO terms to each gene. For GO enrichment analysis, orthogroups were first defined using OrthoFinder (Emms and Kelly 2019). A total of 315 orthogroups containing gene retained exclusively in *Cuscuta* were identified, comprising 2,211 genes. In contrast, 2,067 orthogroups containing gene retained exclusively in non-Cuscuta species were identified, comprising 9,775 genes. Selected genes sets were tested for overrepresentation of GO terms using the clusterProfiler R package (Wu et al. 2021), and statistically significant GO terms were identified based on adjusted *p*-values (Benjamini-Hochberg correction).

## Funding

This work is supported by Suntory Holdings Ltd.

## Acknowledgements

We thank Dr. Tatsuya Wakasugi and Dr. Masayuki Yamamoto (University of Toyama), Koki Nagao (Okayama University), Dr. Kazuya Ishikawa (Ritsumeikan University), Teppei Shinke (Suntory Flowers Ltd.), Dr. Petra Světlíková (OIST), Dr. Hideyuki Arie, Dr. Yoshihide Matsuo and Kazunori Kitao (Suntory Global Innovation Center Ltd.), Dr. Akito Nishizawa (GeneBay, Inc.), Dr. Josef Patzak and Dr. Karel Krofta (Hop Research Institute Co. Ltd.), Dr. Jiří Macas (Biology Centre, Czech Academy of Sciences), Dr. Jun Murata and Dr. Manabu Horikawa (Suntory Foundation for Life Sciences), Dr. Hiroyuki Tanaka (Institute of Science Tokyo), Dr. Atsushi Hoshino (National Institute for Basic Biology), Dr. Koh Aoki (Osaka Metropolitan University) and Dr. Masayuki Maki (Botanical gardens, Tohoku University) for their valuable discussions, technical support, and helpful comments on this study.

## Author Contributions

Tenta Segawa: plant collection, genome analysis, and manuscript writing.

Shima Yoshizumi and Hiromi Toyonaga: molecular biology experiments.

Akira Shiraishi: GO analysis.

Kyoko Sato: chromosome observation.

Takahiro Yamabe and Takehiko Itoh: genome assembly.

Motoshige Takagi: DNA sequencing.

Ryusuke Yokoyama and Masaki Takagawa: preparation of experimental materials.

Eiichiro Ono: plant collection, project organization, and manuscript writing.

## Disclosures

None declared.

## Data availability

The genome assemblies are available at Plant GARDEN (t267557.G001 and t132261.G001). The raw sequence reads are available at DDBJ under accession numbers DRR668413–DRR668421. Specimens from the same population as those used for the genome assembly have been deposited in the Botanical Gardens of Tohoku University; *C. chinensis* (TUS568368, T. Yamashiro MM25026) and *C. campestris* (TUS568367, T. Segawa MM25027), respectively.

## Notes

### Competing Interest Statement

The authors have declared no competing interest.

