## Supplementary Figures for "Chromosome-scale Genome Assemblies of Two Allopolyploid *Cuscuta* Species Uncover Genomic Signatures of Parasitic Lifestyle and Polyploid Evolution"

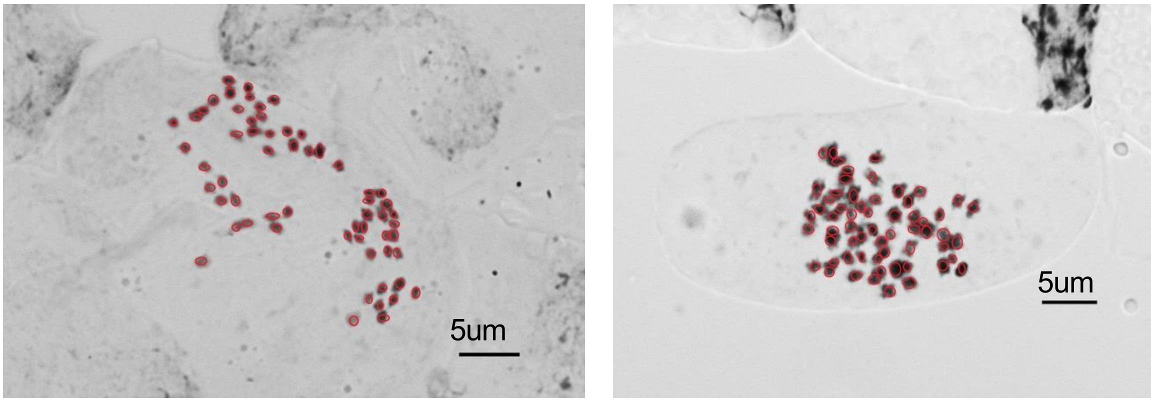


| | | |
|---|---|---|
| *Species* | ***C. campestris*** | ***C. chinensis*** |
| *Karyotype* | 2n=4x=60 | 2n=4x=60 |
| *Genome size* | 514.9 Mbp | 1.50 Gbp |
| *Harvest site (year)* | Kyotango, Kyoto (2024) | Kaibu, Tokushima (2024) |

**Supplementary Figure S1. Chromosome observation with marked boundaries**

Snapshots of chromosome observations corresponding to Fig. 1cd, with the boundaries of each chromosome marked.

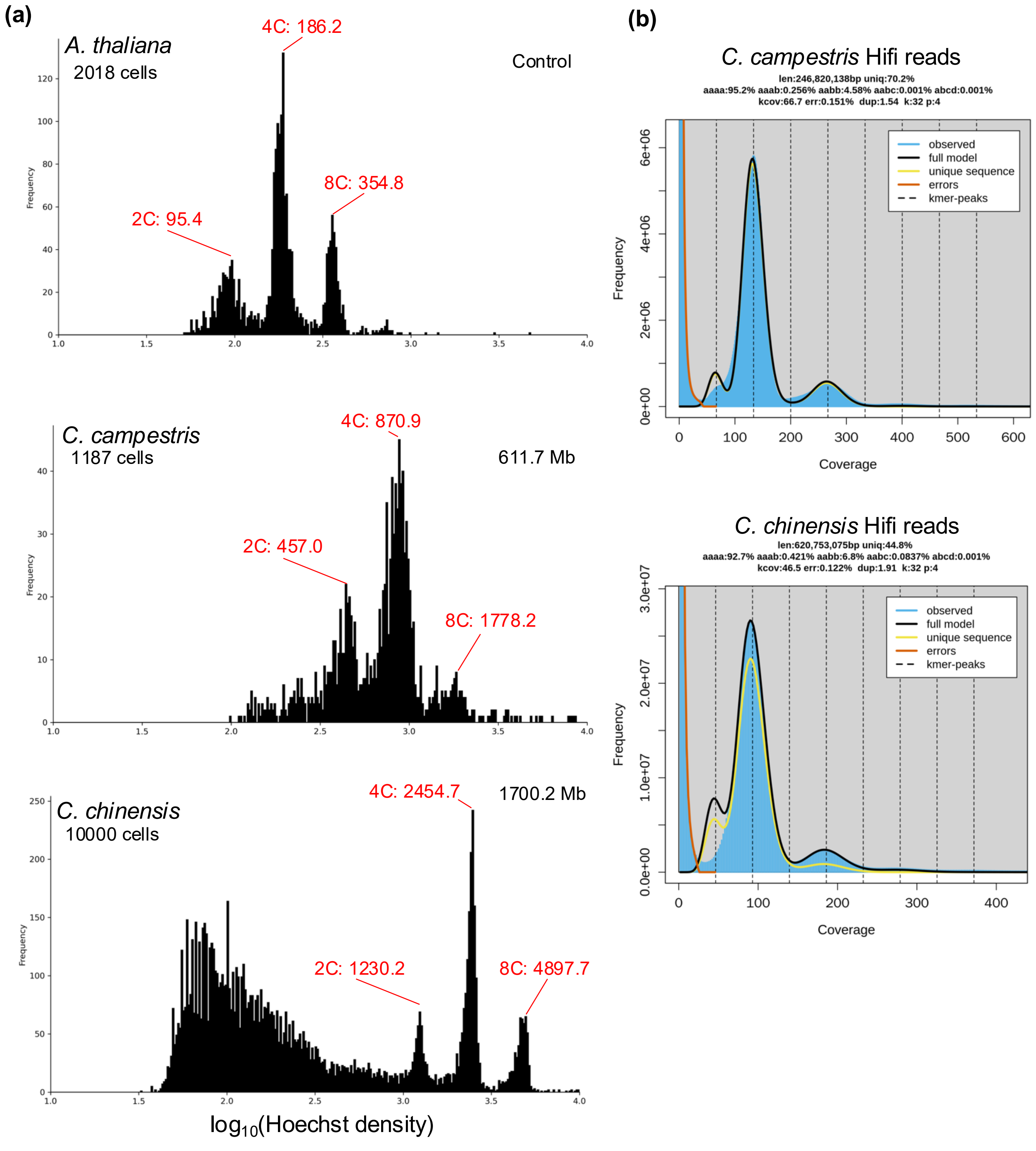


**Supplementary Figure S2. Genome size estimation**

(**a**) Frequency distribution of detection intensities for Hoechst-stained nuclei measured by flow cytometry. (**b**) *K*-mer ($k = 32$) frequency distribution generated from HiFi reads. The predicted size represents the subgenome size, and the actual genome size is expected to be approximately twice this value.

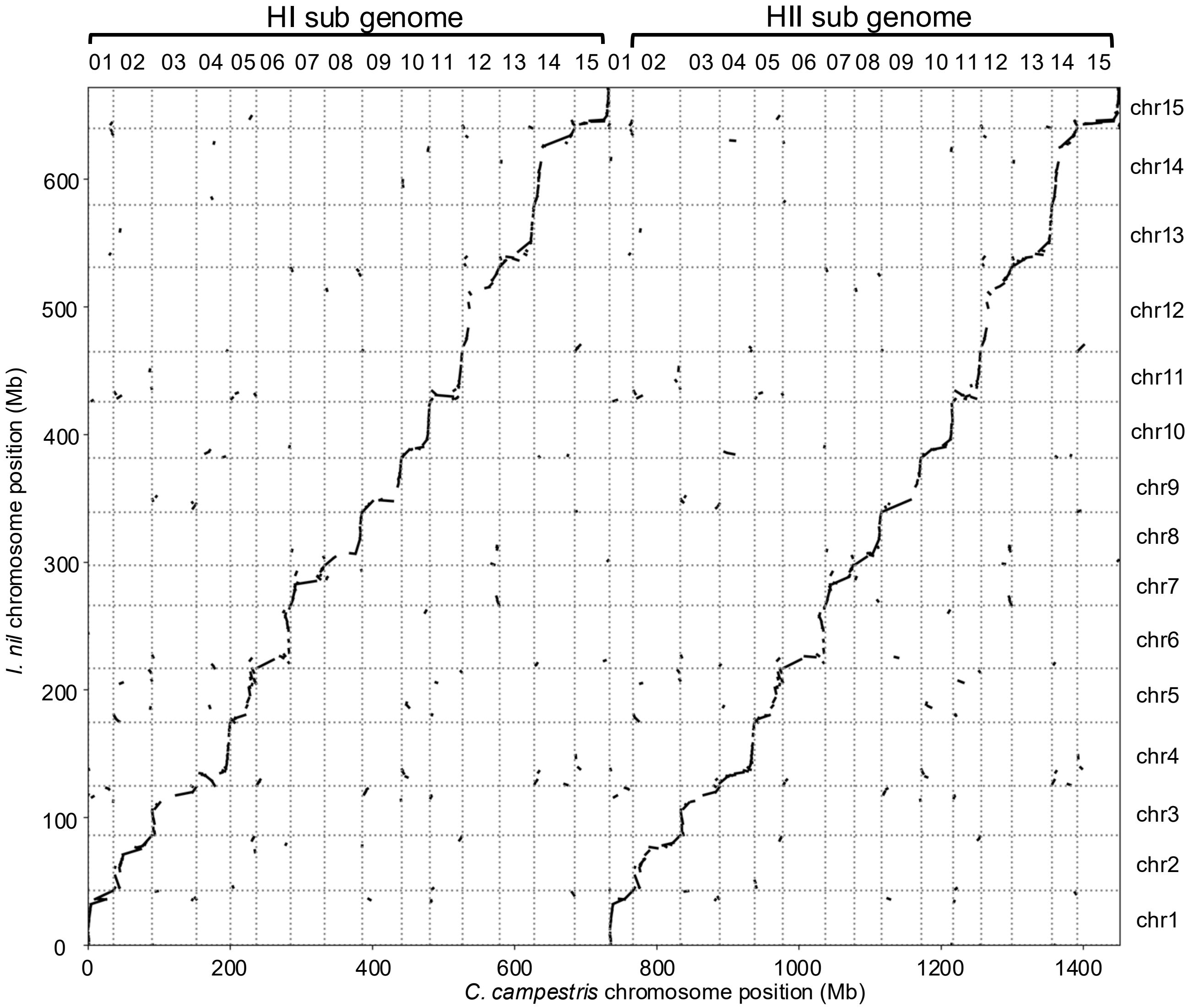


**Supplementary Figure S4. Gene synteny plot between *C. chinesis* and *I. nil.***
Syntenic relationships between *C. chinensis* and *I. nil* are shown.

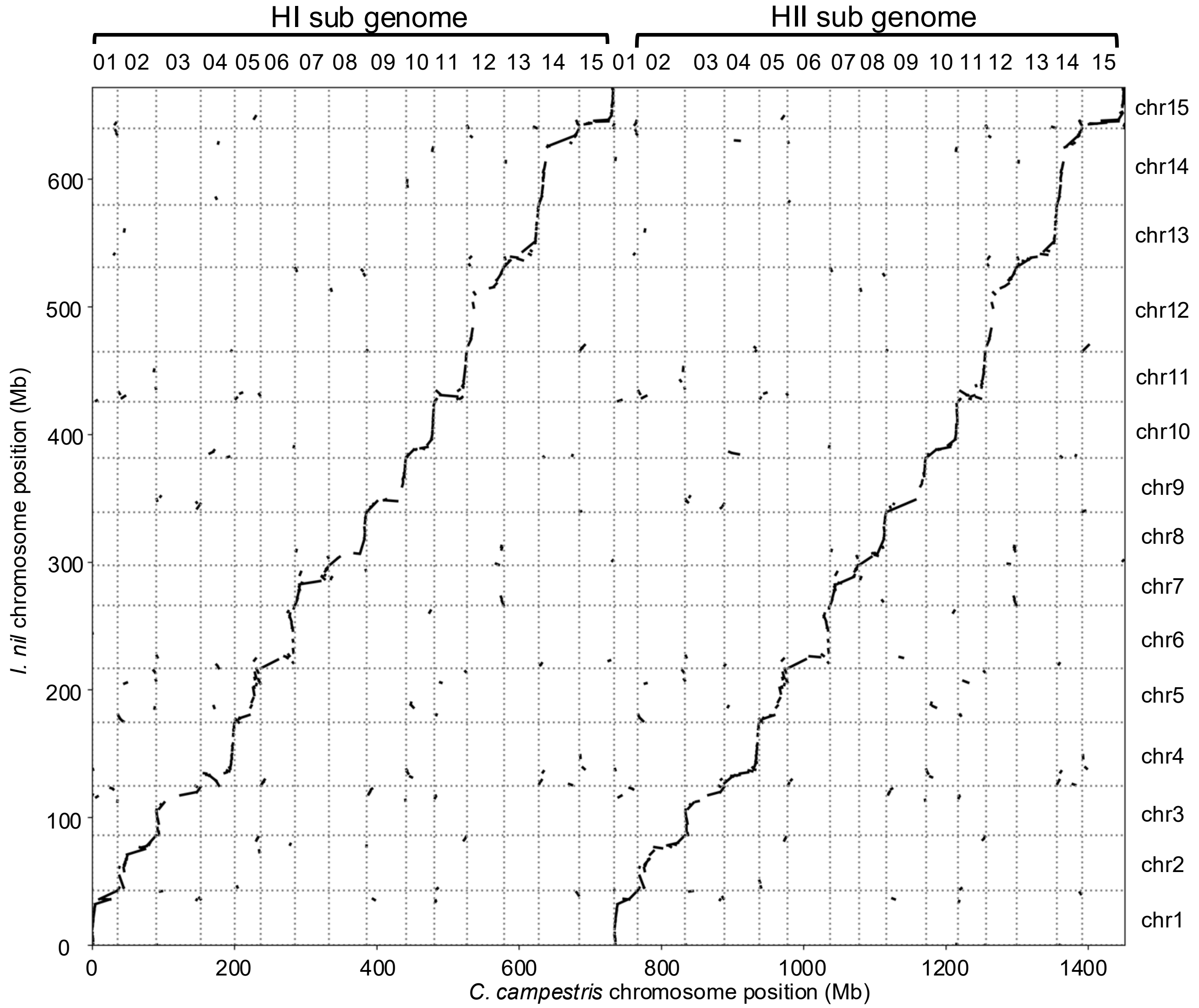


**Supplementary Figure S4. Gene synteny plot between *C. chinesis* and *I. nil.***
Syntenic relationships between *C. chinensis* and *I. nil* are shown.

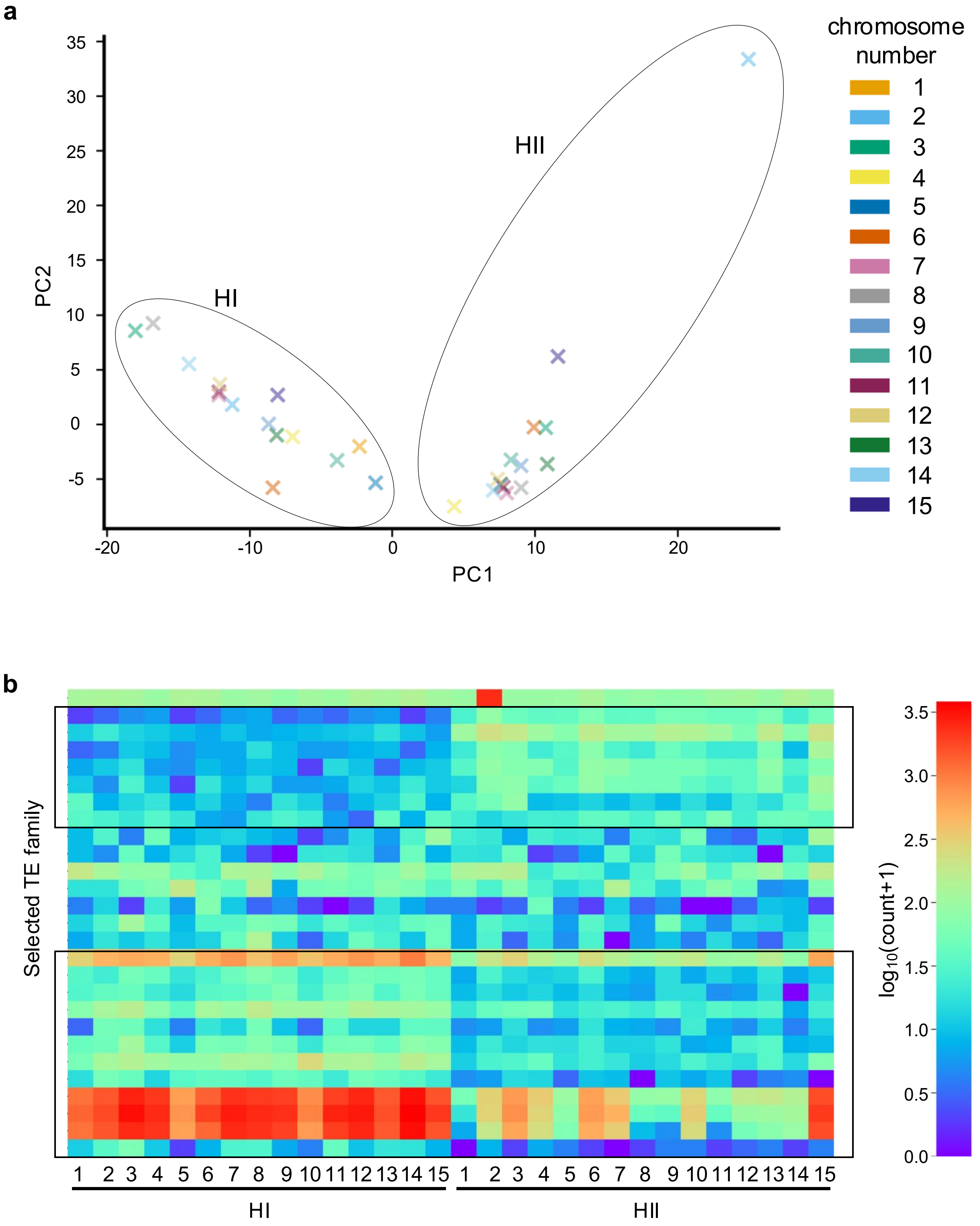


**Supplementary Figure S5. Subgenome-specific TEs in *C. chinensis*.**
(**a**) PCA plot based on the copy numbers of TE families enriched in one of the subgenomes. Chromosome pairs are shown in the same color, and the subgenomes are separated along PC1.
(**b**) Heatmap showing the copy numbers of TE families across chromosomes, based on those used in the PCA. Boxes indicate TE families with significant differences between the HI and HII subgenomes ($p < 0.01$, *t*-test).

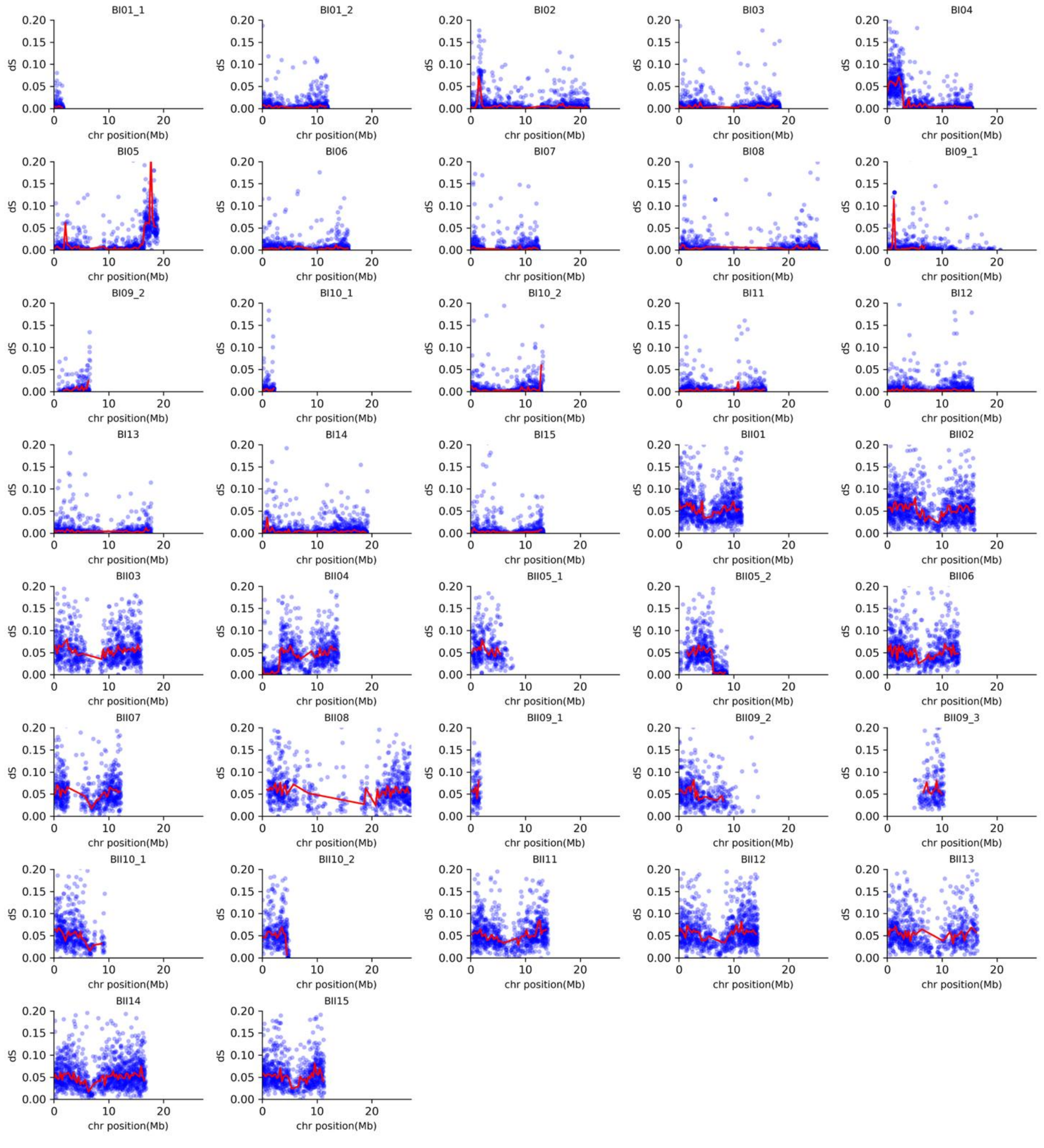


**Supplementary Figure S6. The $d_S$ values of *C. campestris* and *C. australis* genes.**

The blue plots show the $d_S$ values with *C. australis* genes at each gene position. The red lines indicate the median $d_S$ value between *C. campestris* and *C. australis* of 300 kb bins at each chromosomal position.

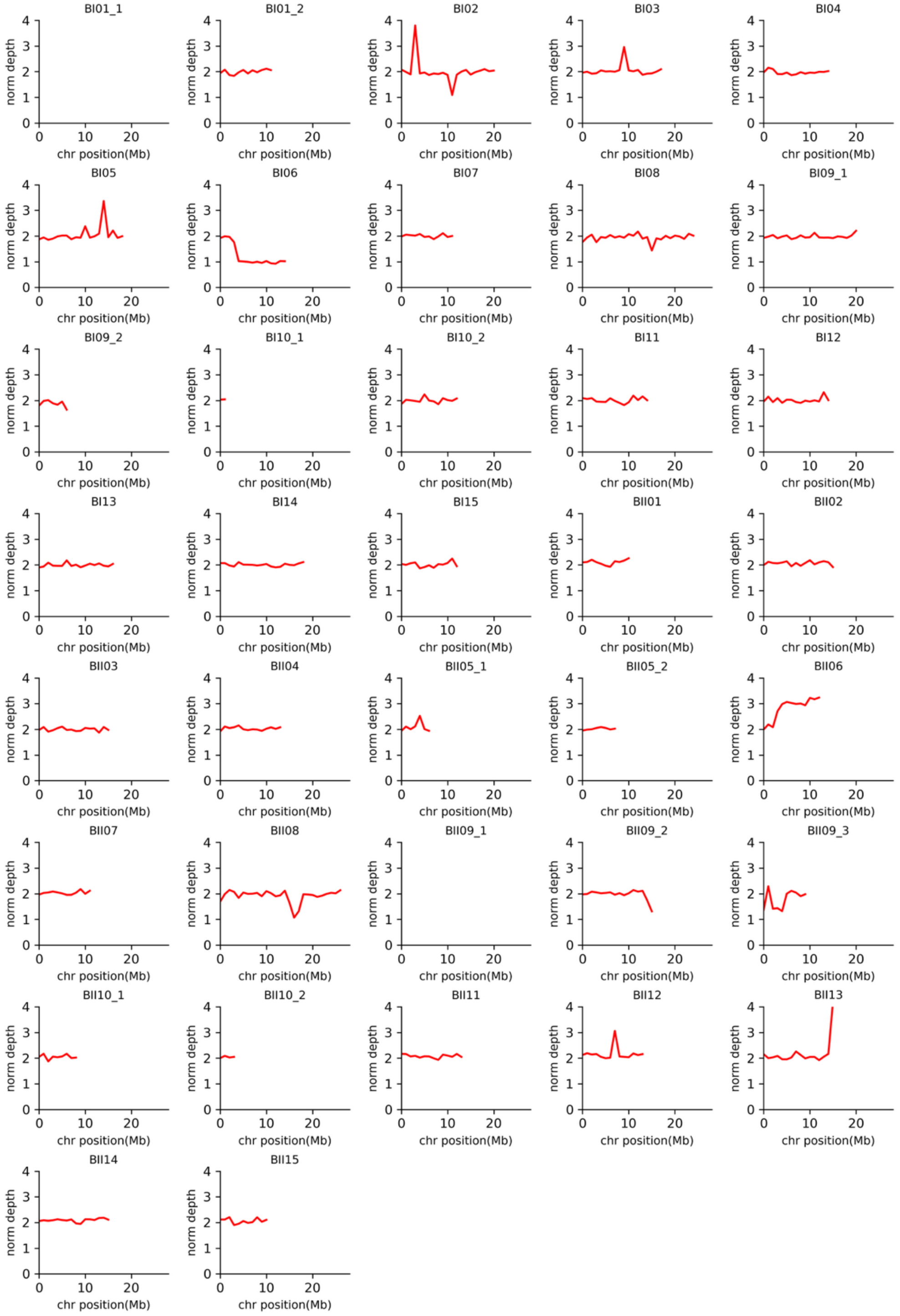


**Supplementary Figure S7. Normalized depth in *C. campestris* self alignment.**
The red line shows the normalized self-aligned depth of 1Mb bins in each chromosome position. Normalized depth is a representation of genome dosage.

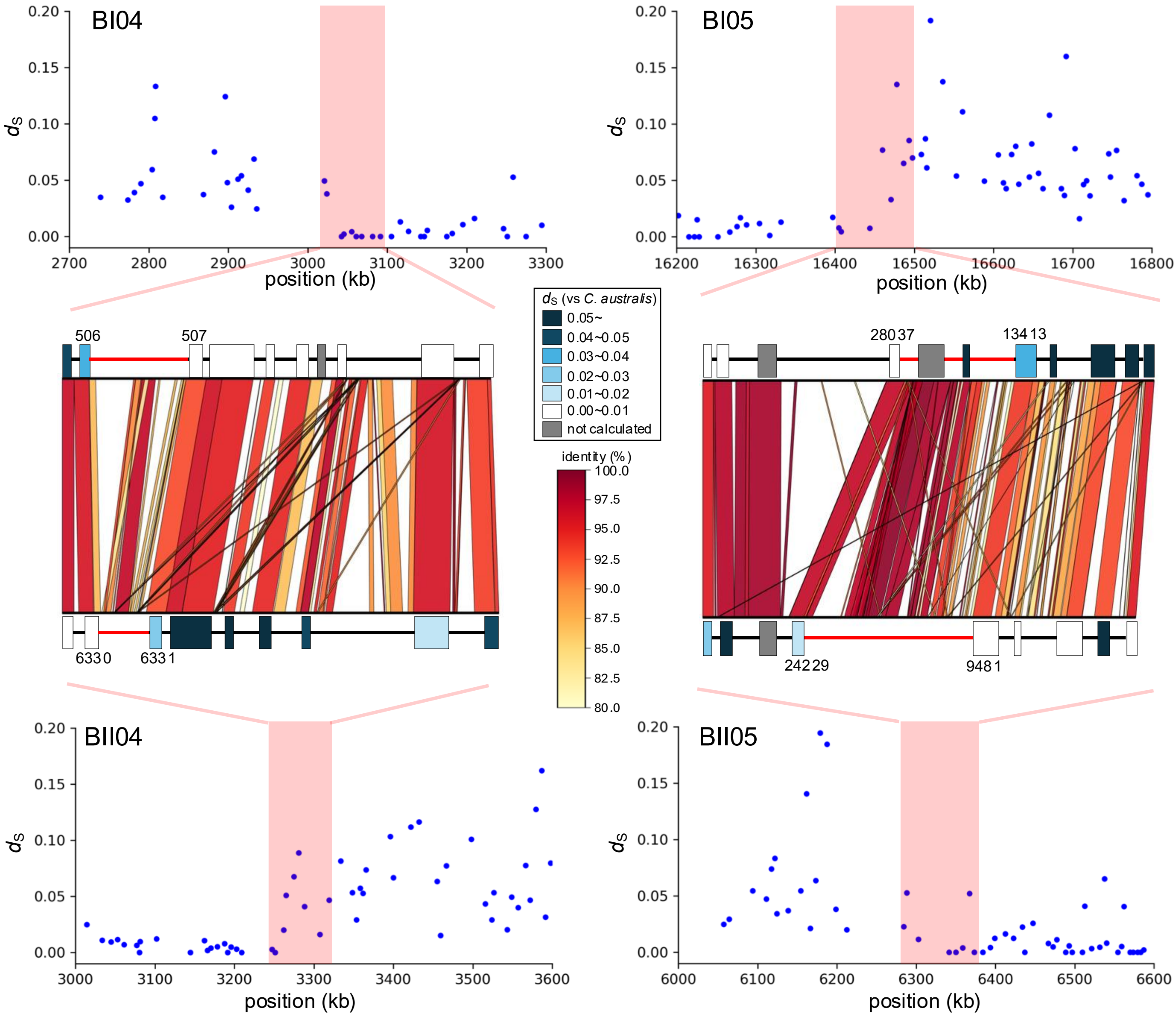


**Supplementary Figure S8. Detailed localization of reciprocal HR regions.**
The upper and lower panels show 600-kb magnified views of the reciprocal HR regions indicated in Figure 3a. Blue dots represent $d_S$ between *C. campestris* and *C. australis* for each gene. Red shading highlights HR regions with noticeable shifts in $d_S$ values. The central panel displays gene positions, their corresponding $d_S$ values, and regions of sequence homology between the subgenomes. Colored boxes indicate gene positions, with colors corresponding to their $d_S$ values. Gray boxes represent genes for which no homolog was detected in *C. australis*. Homologous regions identified via BLASTn are connected and colored according to sequence identity. Genes connected across homologous regions are shown as homoeologous gene pairs.

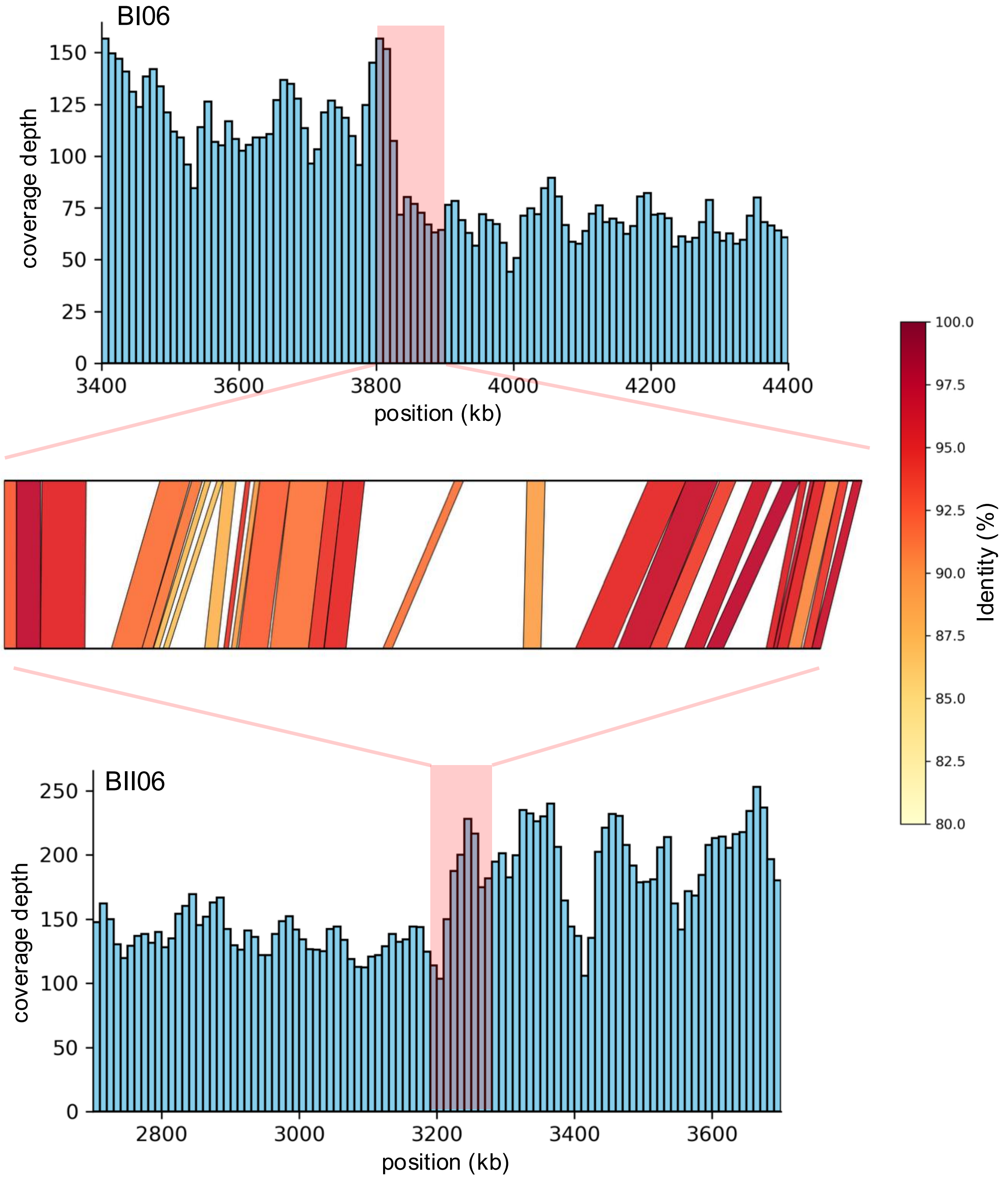


**Supplementary Figure S9. Detailed analysis of coverage depth in non-reciprocal HR regions.** The upper and lower panels show magnified views of the non-reciprocal HR regions presented in Figure 3b. Coverage depth is shown in 10-kb bins across each region. Red highlights indicate regions with pronounced changes in coverage depth. The central panel displays homologous regions between subgenomes within the highlighted areas. Homoeologous region identified via BLASTn are colored based on sequence identity.

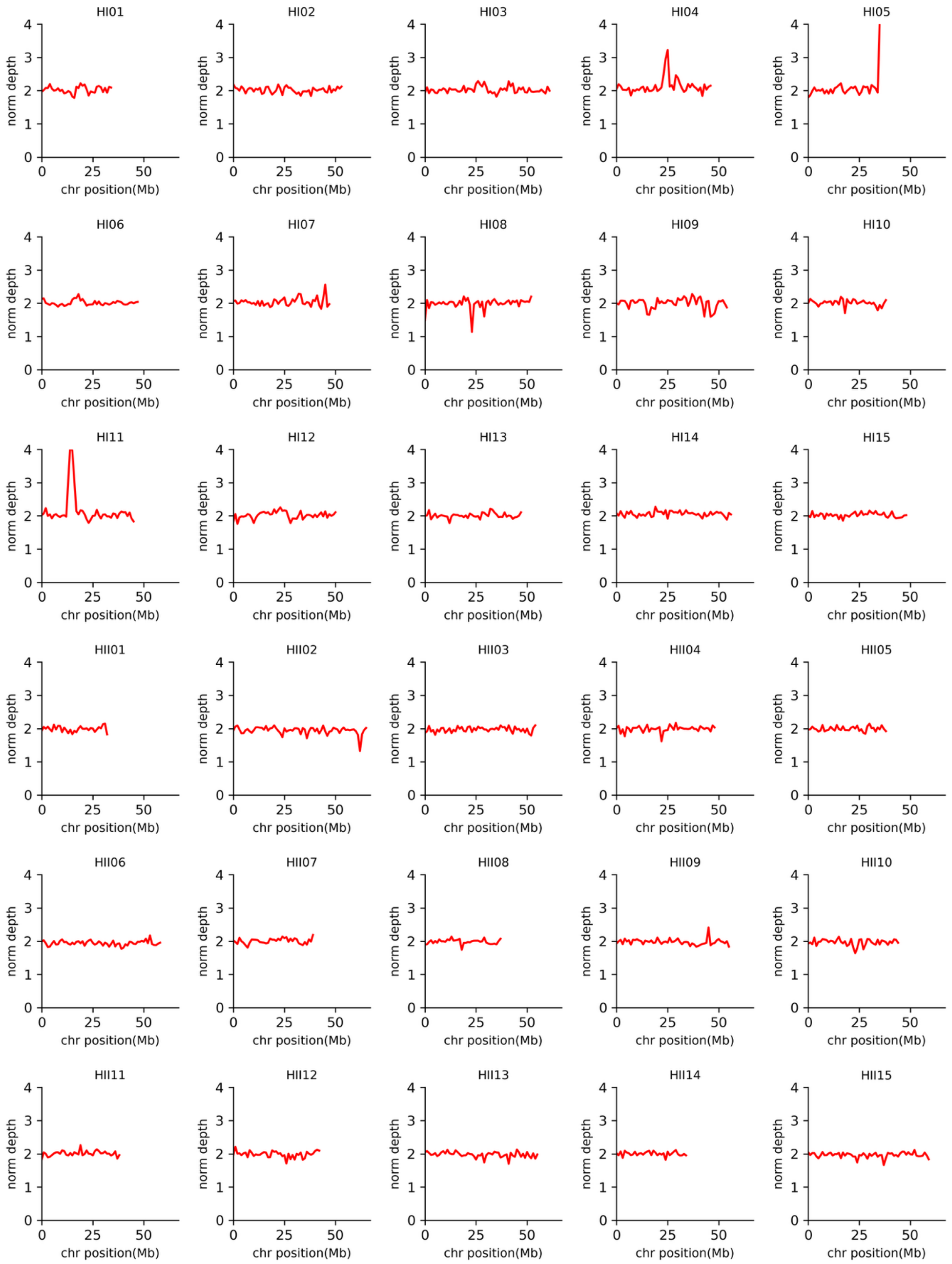


**Supplementary Figure S10. Normalized depth in *C. chinensis* self alignment.**
The red line shows the normalized self-aligned depth of 1Mb bins in each chromosome position. Normalized depth is a representation of genome dosage.

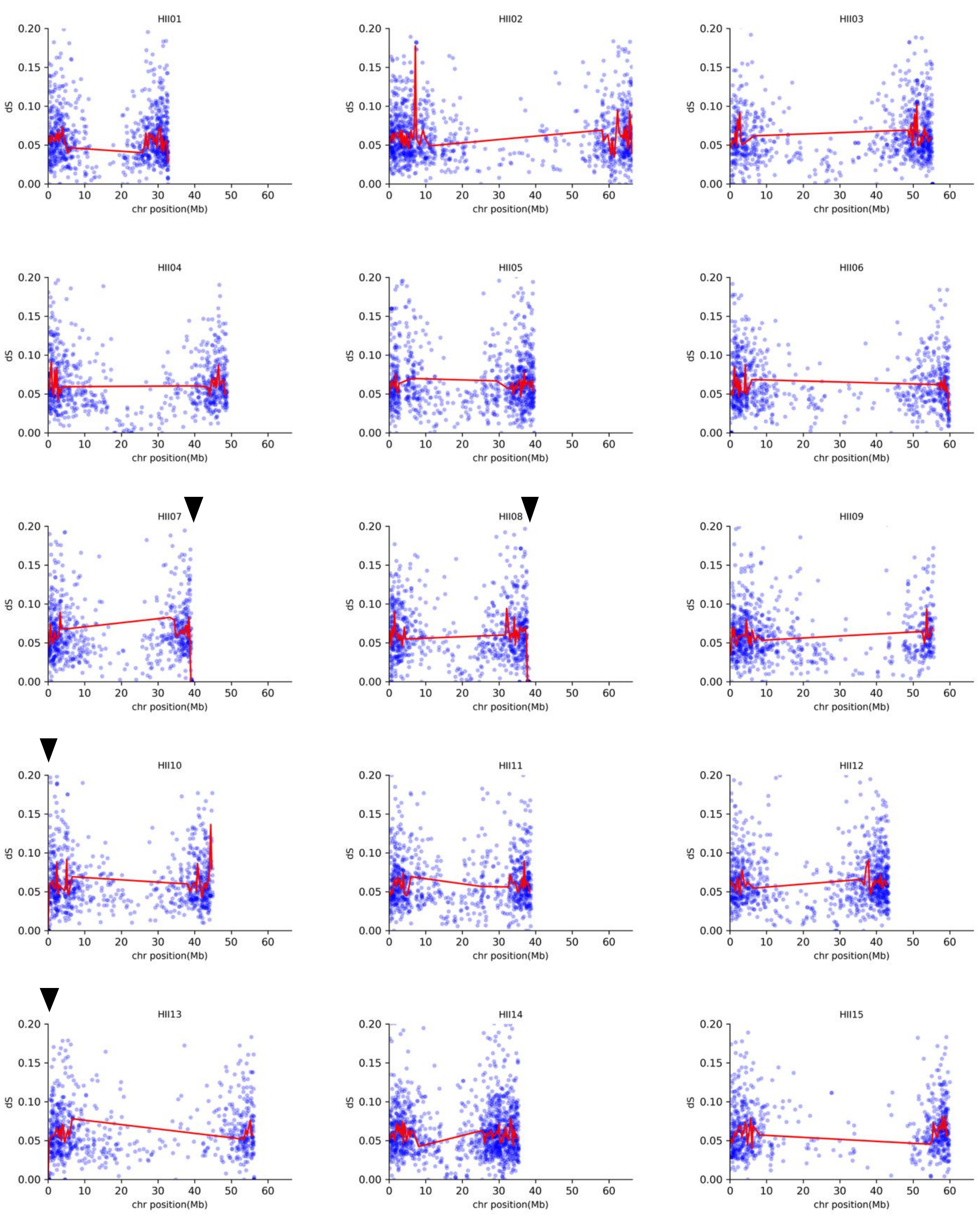


**Supplementary Figure S11. The $d_S$ values of *C. chinensis* genes between subgenome HI and HII.** The blue plots show the $d_S$ values with *C. chinensis* genes at each HII gene position. The red lines indicate the median $d_S$ value between HI and HII of 300 kb bins at each chromosomal position. Black arrowheads indicate non-reciprocal HR candidate sites.

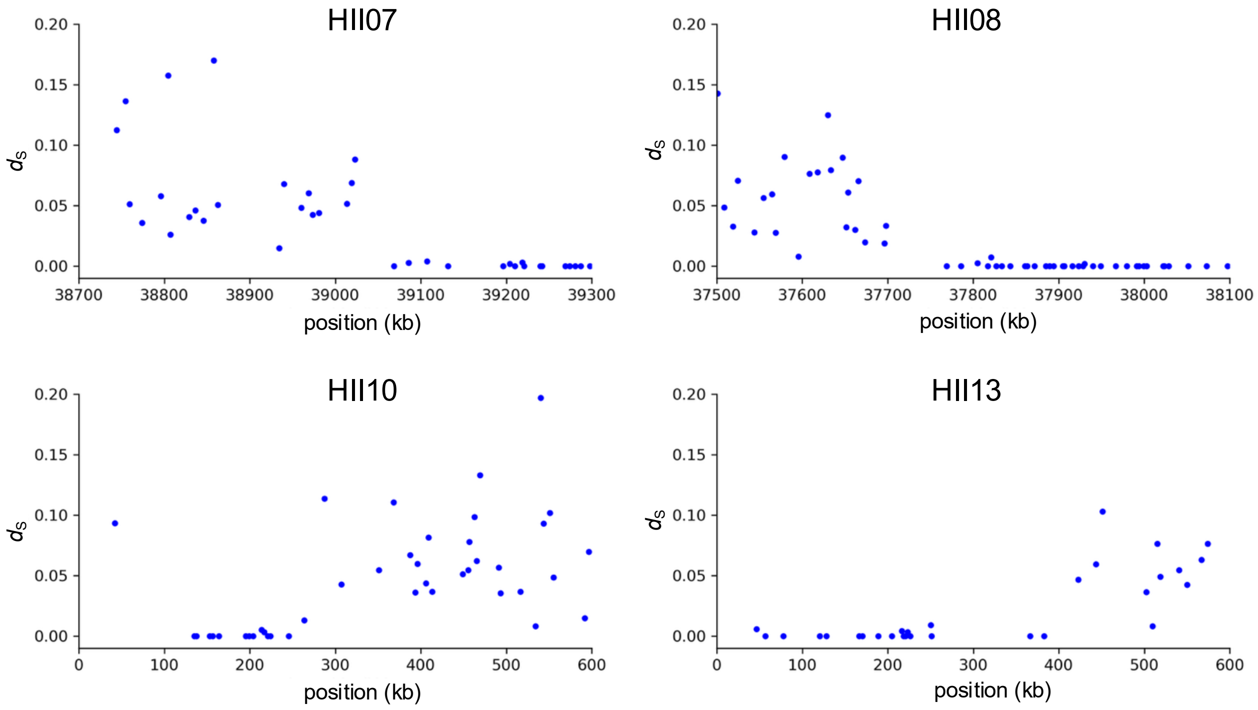


**Supplementary Figure S12. Detailed analysis of non-reciprocal HR sites in *C. chinensis*.**
Each panel shows a magnified view of the non-reciprocal HR sites identified in Supplementary Fig. S11. Blue plots represent the $d_S$ values between the HI and HII subgenomes for each gene. Genomic positions are shown relative to the HII subgenome.

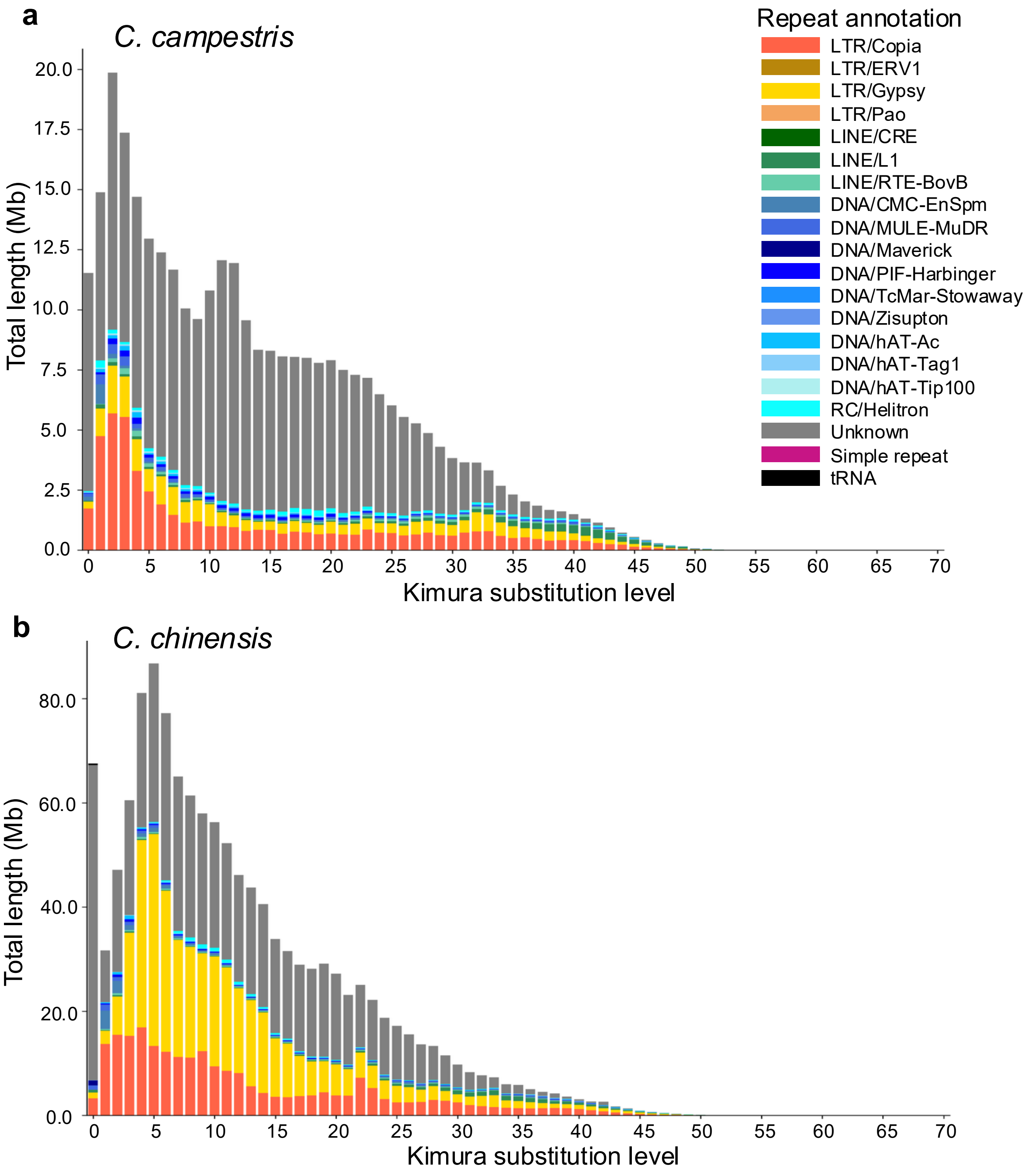


**Supplementary Figure S13. Kimura substitution level of repeat elements in *C. campestris* and *C. chinensis.***

Stacked bar plots show the frequency distribution of Kimura substitution levels for each repeat element in *C. campestris* (**a**) and *C. chinensis* (**b**). Colored bars represent different repeat element families identified by RepeatMasker.

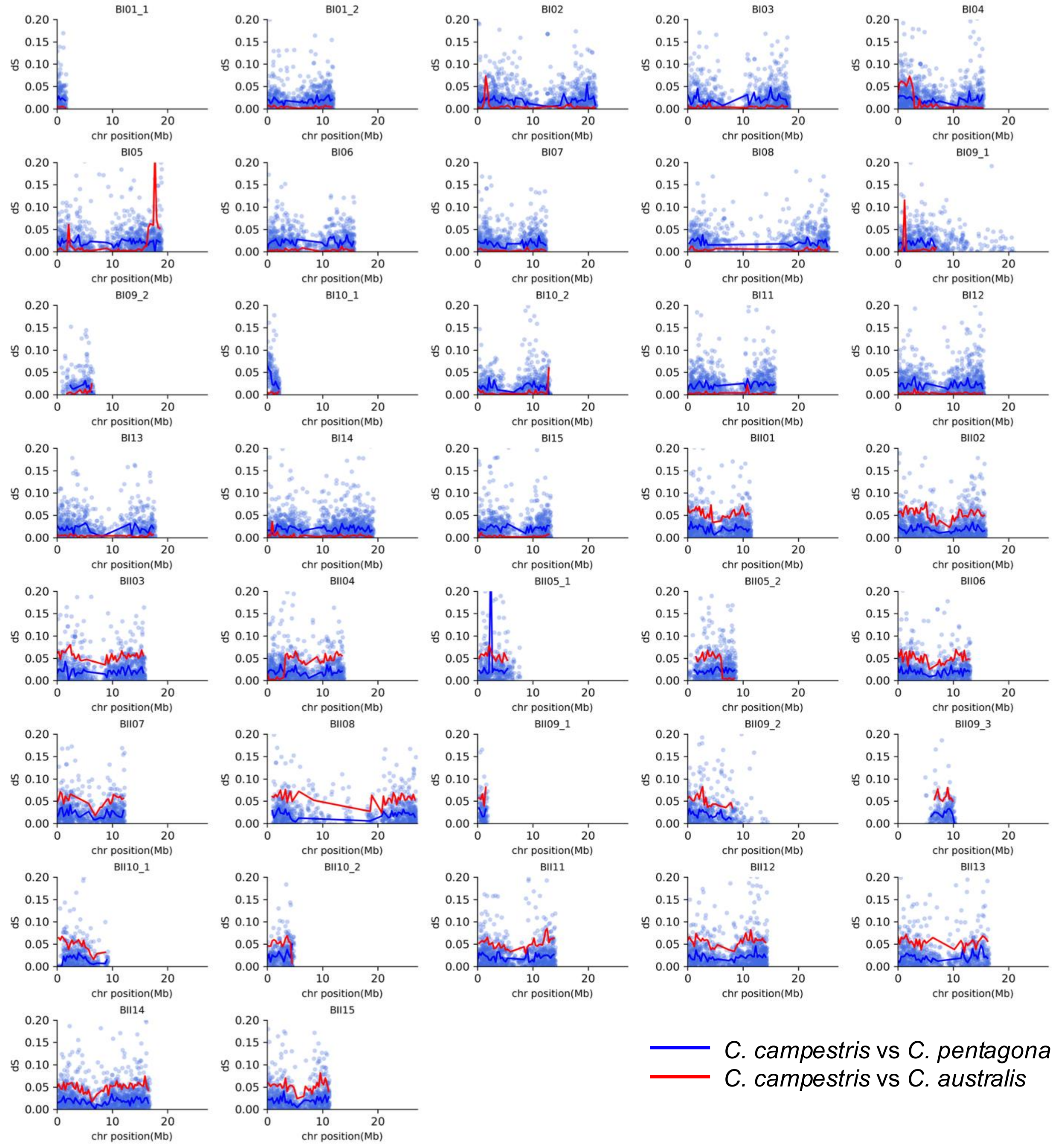


**Supplementary Figure S14. The $d_S$ values of *C. campestris* and *C. pentagona* genes.**

The blue plots show the $d_S$ values with *C. pentagona* genes at each gene position of *C. campestris*. The blue and lines indicate the median $d_S$ value compared with *C. pentagona* and *C. australis*, respectively of 300 kb bins at each chromosomal position.

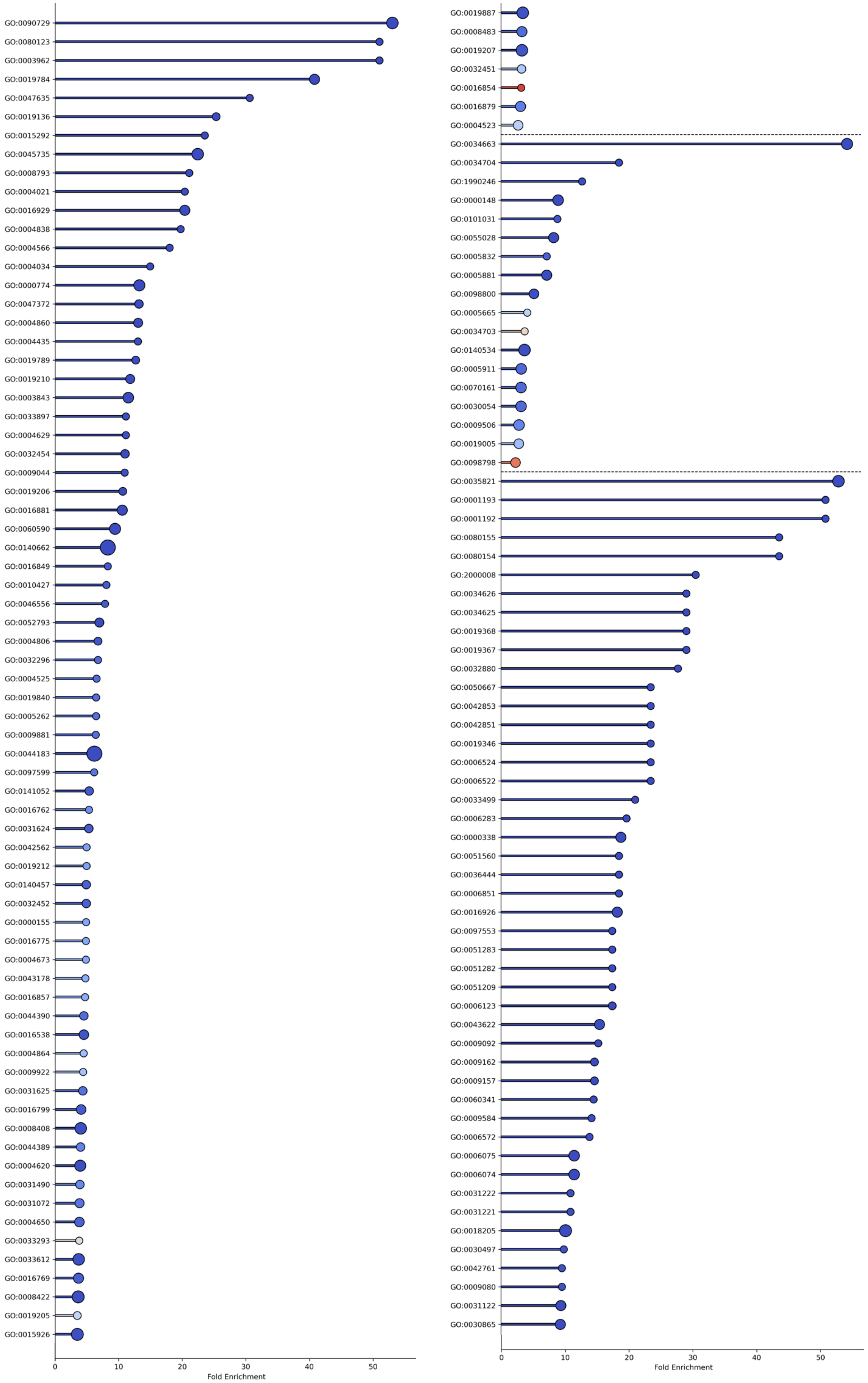


**Supplementary Figure S15. Continued…**

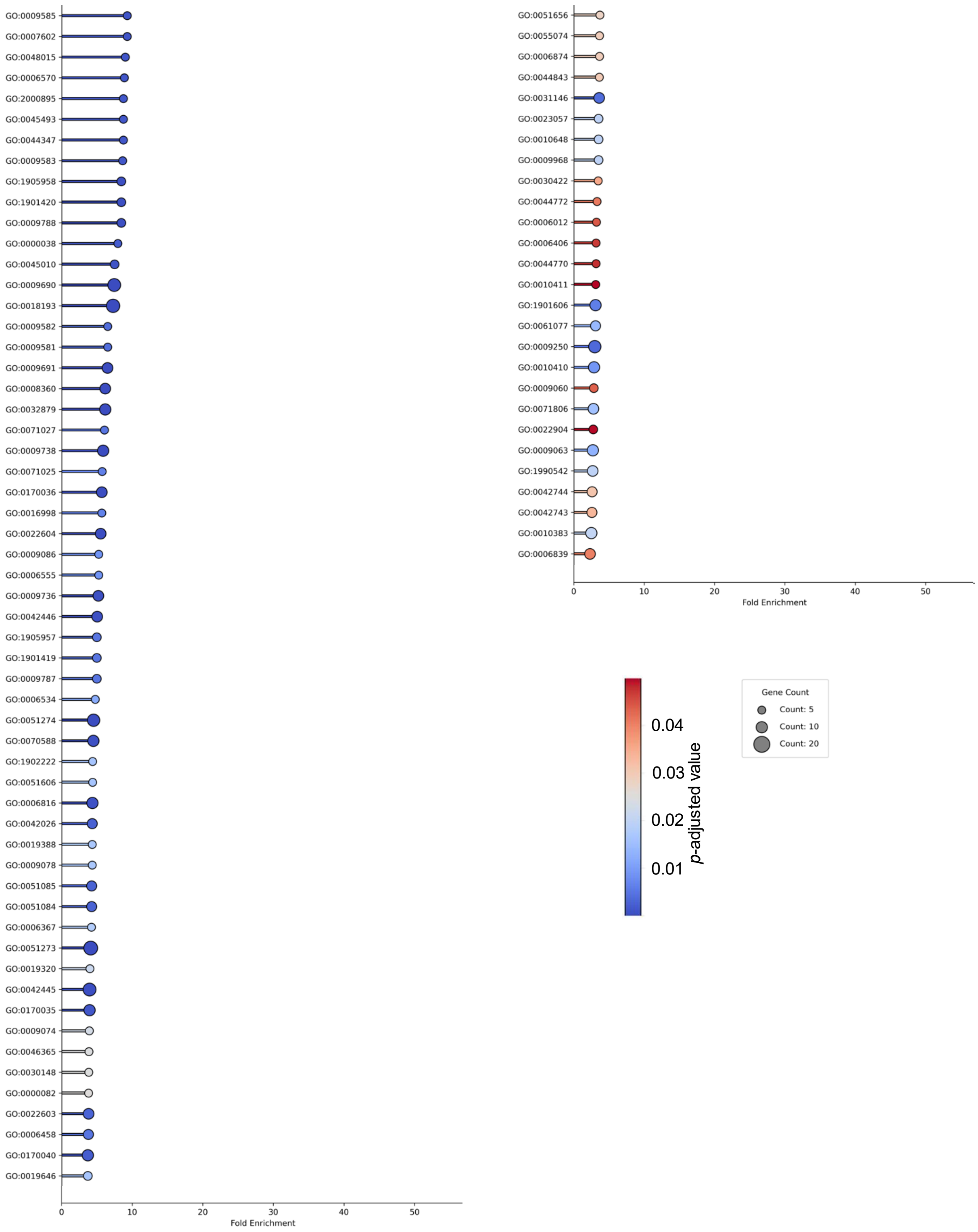


**Supplementary Figure S15. GO enrichment of genes gained in the genus *Cuscuta*.**
GO enrichment analysis of orthogroups specifically gained in *Cuscuta*. Details of each enriched GO term are provided in Supplementary Table S8.

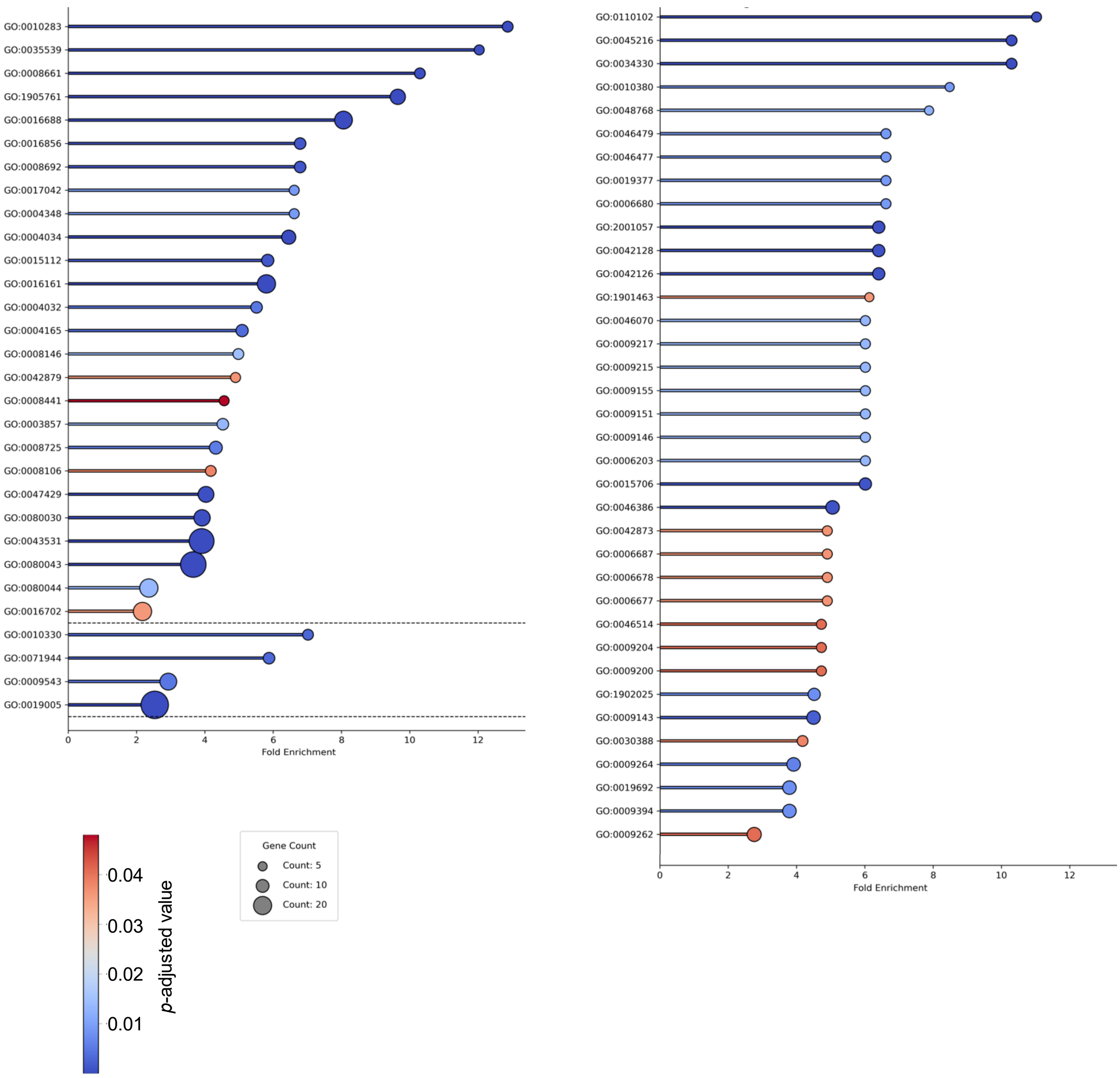


**Supplementary Figure S16. GO enrichment of genes lost in the genus *Cuscuta*.**
GO enrichment analysis of orthogroups specifically lost in *Cuscuta*. Details of each enriched GO term are provided in Supplementary Table S7.
